# Highly accurate reference and method selection for universal cross-dataset cell type annotation with CAMUS

**DOI:** 10.1101/2025.01.13.632744

**Authors:** Qunlun Shen, Shuqin Zhang, Shihua Zhang

## Abstract

Cell type annotation is a critical and essential task in single-cell data analysis. Various reference-based methods have provided rapid annotation for diverse single-cell data. However, how to select the optimal references and methods is often overlooked. To this end, we present a cross-dataset cell-type annotation methodology with a universal reference data and method selection strategy (CAMUS) to achieve highly accurate and efficient annotations. We demonstrate the advantages of CAMUS by conducting comprehensive analyses on 672 pairs of cross-species scRNA-seq datasets. The annotation results with references selected by CAMUS achieved substantial accuracy gains (25.0-124.7%) over random selection strategies across five reference-based methods. CAMUS achieved high accuracy in choosing the best reference-method pair among 3360 pairs (49.1%). Moreover, CAMUS showed high accuracy in selecting the best methods on the 80 scST datasets (82.5%) and five scATAC-seq datasets (100.0%), illustrating its universal applicability. In addition, we utilized the CAMUS score with other metrics to predict the annotation accuracy, providing direct guidance on whether to accept current annotation results.

## Introduction

Single-cell technologies such as single-cell RNA sequencing (scRNA-seq) (Kharchenko 2021), single-cell spatial transcriptomics (scST) (Longo et al. 2021), and single-cell Assay for Transposase-Accessible Chromatin using sequencing (scATAC-seq) (Fang et al. 2021) have revolutionized our understanding of cellular diversity and complexity. These technologies enable researchers to investigate the transcriptomic, spatial, and epigenomic landscapes at single-cell resolution, offering unprecedented insights into understanding organismal development, physiological responses, and disease progression (Jia et al. 2018; Longo et al. 2021; Srivatsan et al. 2021).

One critical task in single-cell data analysis is cell type annotation (or cell typing) (Clarke et al. 2021; Brbić et al. 2022), allowing researchers to decipher cell heterogeneity and their functions within the tissue microenvironment. The prevalent manual annotation approach groups single cells into several clusters and annotates them based on unique molecular signatures (Clarke et al. 2021). This process needs prior biological knowledge and is time-consuming. However, as scRNA-seq can profile all genes (typically numbering in the tens of thousands), the clustering results from well-profiled scRNA-seq data are relatively distinct, resulting in high accuracy of cell type annotation. Recently, as extensive scRNA-seq datasets are generated and annotated, reference-based annotation methods have gained prominence (Clarke et al. 2021). These methods effectively transfer cell labels from the reference dataset to the query one by leveraging the well-annotated scRNA-seq datasets. Reference-based methods such as CAME (Liu et al. 2023b), Seurat (Hao et al. 2024), SciBet (Li et al. 2020), scmap (Kiselev et al. 2018), and SCN (Tan and Cahan 2019) have demonstrated rapid cell annotation capabilities without needing prior biological knowledge. These methods potentially enable the cell type identification that may be overlooked or difficult to ascertain through manual annotation and have been successfully extended to various scenarios, encompassing different conditions, technologies, and species (Liu et al. 2023b; Hao et al. 2024).

Cell type annotation for scST data presents even more challenges, as spatial technologies such as seqFISH (Shah et al. 2018), STARmap (Wang et al. 2018), and MERFISH (Zhang et al. 2021) typically profile only hundreds of genes. This limitation results in blurred cluster boundaries and difficulties in manual annotation (Shen et al. 2025). Therefore, most scST studies utilize scRNA-seq datasets as a reference to help annotate scST datasets. For example, scANVI was employed to annotate MERFISH data derived from the kidney and liver (Xu et al. 2021; Liu et al. 2023a); RCTD was used on Slide-seq v2 data collected from the cerebellum and hippocampus (Cable et al. 2022); Tangram was applied to STARmap and MERFISH data originating from the cortex (Biancalani et al. 2021); and STAMapper was evaluated under scST datasets collected from eight scST technologies and five tissues (Shen et al. 2025).

Regarding scATAC-seq data, its extreme sparsity often constrains the performance of cell type annotation (Lin et al. 2022). This limitation necessitates the development of more sophisticated reference-based computational methods to assist cell typing. For instance, GLUE (Cao and Gao 2022), scJoint (Lin et al. 2022), and scDART (Zhang et al. 2022) learn a joint cell embedding from scRNA-seq and scATAC-seq data, which aids in annotating the scATAC-seq data.

Although we have well-designed reference-based cell type annotation methods for various omics datasets, no single method consistently achieves optimal performance across all omics types. Furthermore, for any given tissue or disease, numerous well-annotated scRNA-seq datasets are available as references. The choice of reference can significantly impact the annotation results. For single-cell assays, some cell types may be lost, underrepresented, or annotated incorrectly (Garmire et al. 2024). Naturally, reference and method selection are critical considerations. Without a solid understanding of biological context, users cannot assess the accuracy of algorithmic annotations. Estimating the accuracy of these annotations can also guide users directly in deciding whether to accept a particular annotation. However, to the best of our knowledge, there are currently no specific reference selection tools or method-agnostic optimizers designed for single-cell omics analysis. While several annotation tools perform method-specific tuning, they do not provide a general-purpose framework that can assess and compare multiple reference-method pairs in a consistent and data-driven manner.

To this end, we present a cross-dataset annotation methodology with a universal reference data and method selection strategy (CAMUS) for reference-based single-cell annotation methods. CAMUS prioritizes the annotation performance of reference-based methods by comparing the concordance between the annotation results and the per-clustered labels based on adjusted mutual information (AMI), we refer to this metric as the CAMUS score.

## Results

### Overview of CAMUS

CAMUS aims to prioritize reference-method pairs and select the optimal one, ensuring the highest accuracy for a reference-based single-cell annotation task. CAMUS utilizes the single-cell expression profiles from the query data and the annotation results of various reference-method pairs as inputs (**Figure 1a, STAR Methods**). It initially pre-clusters cells from query data into several distinct clusters (**Figure 1a**) to approximate the cell type composition of the query data. It compares the concordance score between the pre-clustering and annotation results with various methods and references. A higher CAMUS score indicates a potentially higher annotation accuracy (**STAR Methods**). CAMUS can be applied to three distinct cell type annotation scenarios: (1) Cross-species (including species from human, mouse, zebrafish, chick, lizard, turtle, and macaque); (2) Cross modalities (from scRNA-seq dataset to scST dataset); (3) Cross omics (from scRNA-seq dataset to scATAC-seq dataset) (**Figure 1b**). CAMUS effectively prioritizes reference-method pairs with rankings highly correlated with the actual annotation accuracy. Furthermore, CAMUS precisely estimates annotation accuracy by utilizing CAMUS score and additional cellular indices as predictors (**STAR Methods**), offering users reliable guidance (**Figure 1c**).

**Figure 1.**
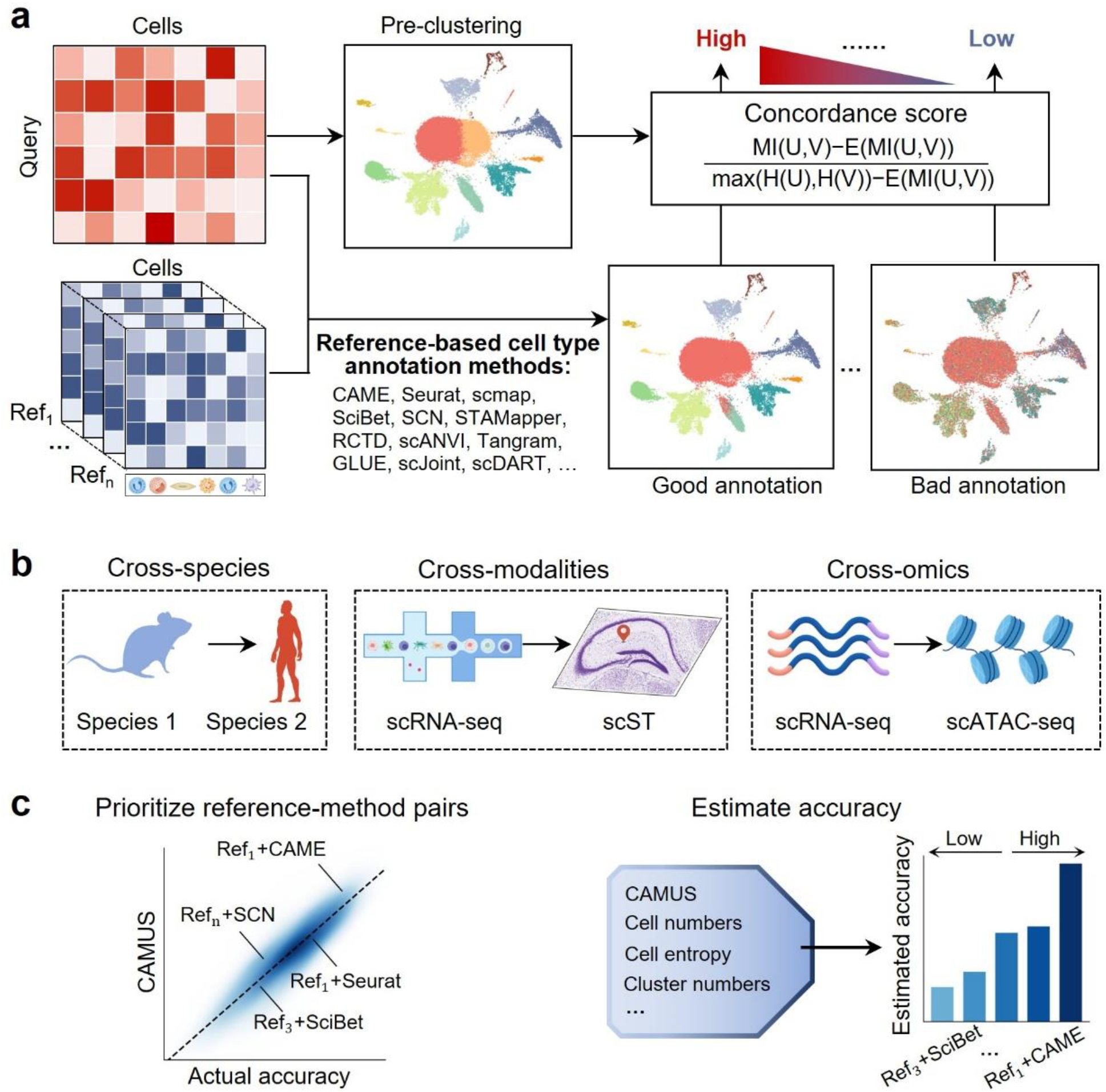
Overview of CAMUS. **a**, CAMUS first pre-clusters single-cell data into several distinct clusters and then combines diverse reference datasets and reference-based annotation methods to annotate the query dataset. CAMUS compares the concordance score between the pre-clustering and the various annotation results. A higher CAMUS score indicates a potentially higher annotation accuracy. **b**, CAMUS is applied to three cell type annotation scenarios: (1) Cross-species (including species from human, mouse, zebrafish, chick, lizard, turtle, and macaque); (2) Cross-modalities (from scRNA-seq dataset to scST dataset); (3) Cross-omics (from scRNA-seq dataset to scATAC-seq dataset). CAMUS helps to define the rank of reference-method pairs, which is positively correlated with the corresponding actual accuracy. The CAMUS score serves as a predictor to estimate annotation accuracy for users.

### CAMUS enables highly accurate reference selection for cross-species annotation

To demonstrate the effectiveness of CAMUS in prioritizing the reference datasets, we gathered 672 pairs of cross-species datasets from seven species and five tissues, each pair accompanied by manual annotations serving as the ground truth (**STAR Methods, Supplemental Table S1)**. There are 53 distinct query datasets, 47 of which have three or more reference datasets. We applied five reference-based annotation methods (i.e., CAME (Liu et al. 2023b), Seurat (Hao et al. 2024), SciBet (Li et al. 2020), scmap (Kiselev et al. 2018), and SCN (Tan and Cahan 2019)) to all the data pairs.

Across all 3,360 data pairs, CAMUS scores demonstrated significant positive correlations with annotation accuracy for the five methods (**Figure 2a**). In ∼99 % of cases, references with higher CAMUS scores produced more accurate annotation results (**Supplemental Fig. S1a**). A closer look at four cross-species retinal datasets (∼30 references each) confirmed that the references ranked first by CAMUS always yield the best (or second-best) annotations, regardless of the annotation method used (**Figure 2b-e**). Specifically, the optimal references selected by CAMUS consistently showed high accuracy, particularly for the better-performing annotation methods CAME and Seurat. Specifically: in the Human Menon microfluidics dataset, CAME (95.55%), Seurat (90.48%), SciBet (76.37%), scmap (82.15%), and SCN (78.90%); in the Mouse NMDA 12hr dataset, CAME (98.0%), Seurat (93.89%), SciBet (87.76%), scmap (85.82%), and SCN (63.86%); in the Chick P10 dataset, CAME (93.23%), Seurat (84.02%), SciBet (73.10%), scmap (69.88%), and SCN (58.32%); and in the Zebrafish LD 4hr dataset, CAME (77.47%), Seurat (61.09%), SciBet (43.63%), scmap (50.22%), and SCN (34.99%). Furthermore, the reference selection has a substantial impact on annotation accuracy. The best reference can improve it by more than 150% compared to the worst (**Figure 2c**), highlighting the importance of reference selection.

**Figure 2.**
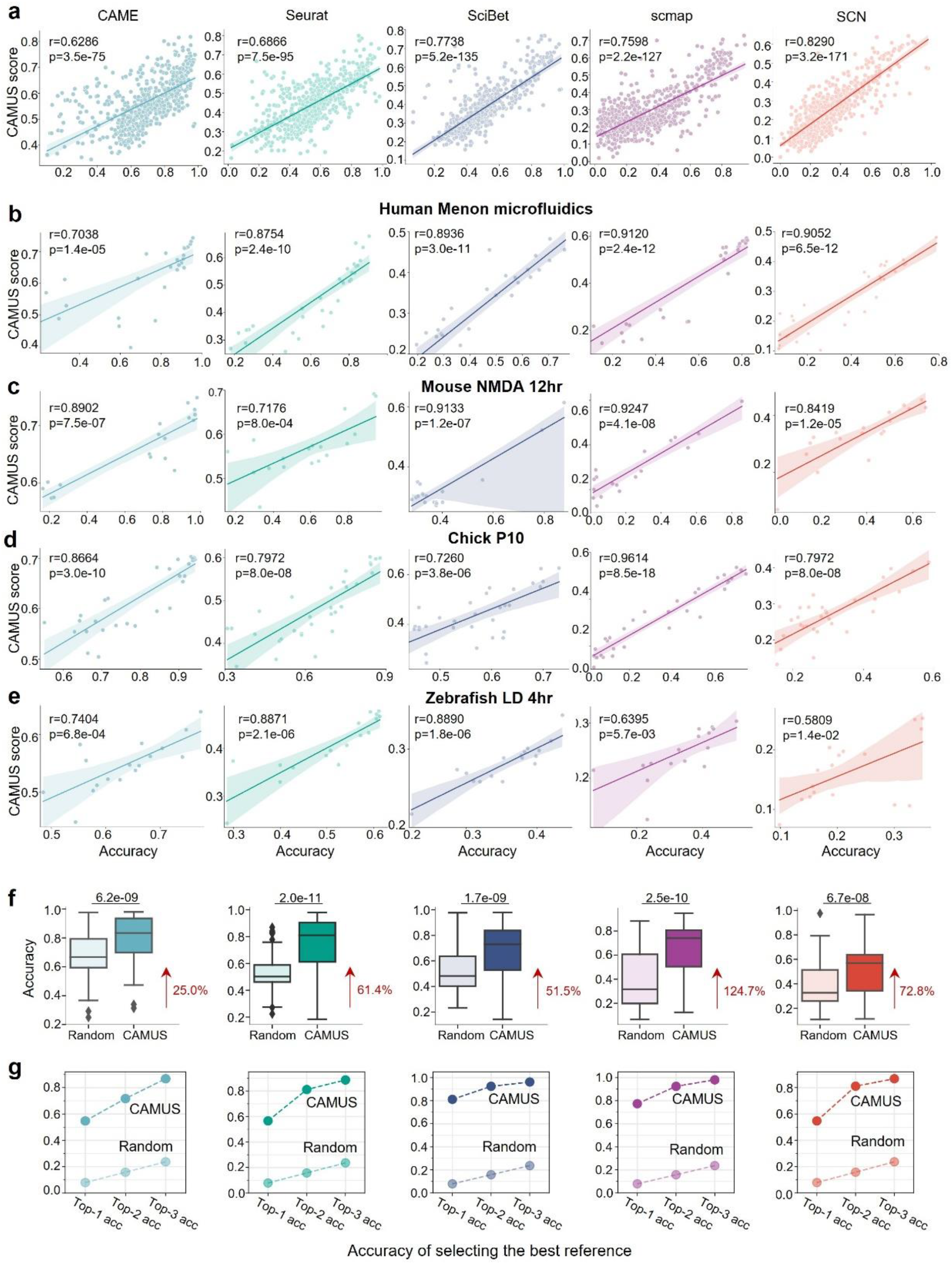
Application of CAMUS for reference selection in cross-species cell type annotation. **a-e**, Scatter plot showing the significant positive correlation between the CAMUS score and annotation accuracy in all cross-species data pairs (n=672) (**a**), and specific query datasets from humans, mice, chicks, and zebrafishes (**b-e**) for five cell annotation methods, i.e., CAME, Seurat, SciBet, scmap, and SCN. **f**, Boxplot indicating the accuracy of cell type annotation. The references are selected by CAMUS or randomly. For the random selection, we performed five independent replicates for each setting and report the average values. The p-value was calculated using the paired t-test. The data following the red arrow represents the percentage improvement for the median value relative to random selection. **g**, Dot plot showing the top-1, top-2, and top-3 accuracy of selecting the best reference by CAMUS or randomly.

Next, we quantitatively evaluated the improvements by selecting references using CAMUS. For five distinct methods, the annotation accuracy improves over random selection strategies, ranging from 25.0% to 124.7% (**Figure 2f**) using the best reference selected by CAMUS compared to random selection for all 672 data pairs. Substantial gains were also observed in both macro F1 score (12.1-99.0%) and weighted F1 score (24.3-127.8%) (**Supplemental Fig. S1b, c**). To quantify the impact of reference-set size on CAMUS, we subsampled the original reference data to 50% and evaluated the accuracy of both CAMUS and random sampling. The performance of CAMUS still significantly outperforms random selection (**Supplemental Fig. S1d**). This implies that reference data selection can enhance annotation accuracy and rare cell type identification. Furthermore, we observed significant variation in the dependence on references among different annotation methods. For instance, under the random selection strategy, CAME achieved significantly higher accuracy than Seurat (median 66.71% vs 50.21%, *p*=2.2e-12). However, this gap was greatly reduced when reference selection was guided by the CAMUS score (median 83.41% vs 81.01%, *p*=7.5e-04), and the annotation accuracy improved significantly (**Figure 2f**). This phenomenon was similarly reflected in other methods. That is to say, prioritizing the reference datasets is essential for more accurate annotation of a given query data. CAMUS significantly outperforms random selection in consistently identifying the top 1-3 references with much higher accuracy (**Figure 2g**). Moreover, CAMUS demonstrates strong robustness in selecting the best reference across different clustering resolutions and various clustering methods and achieved stable top-1 accuracy in selecting the best reference (∼55% for CAME, Seurat, and SCN, ∼80% for SciBet and scmap) (**Supplemental Fig. S2a, b**). Additionally, we divided the query datasets into those with fewer than 11 cell types (n=23) and those with more than 11 cell types (n=31). CAMUS consistently maintained robust performance across varying numbers of cell types, as well as under different clustering strategies and parameter settings (**Supplemental Fig. S1c**).

In single-cell data, when the number of the annotated cell types and the pre-clustering differ substantially (e.g., the reference-based annotation methods annotate cell subtypes while the pre-clustering only recovers major cell types), we want the CAMUS score to remain relatively high. We provide a formal theoretical justification for the superiority of AMI over Adjusted Rand Index (ARI) and Fowlkes-Mallows Index (FMI) (**Supplemental Note 2**). Therefore, we choose AMI as part of the CAMUS framework. The real-world experiments on cross-species tasks further confirmed these results, where the correlation between AMI and the accuracy consistently outperformed that of FMI and ARI across all five methods (**Supplemental Fig. S3**).

### CAMUS enables highly accurate method selection for cross-species annotation

For all the 672 pairs of cross-species datasets considered, the average correlation between the CAMUS score and the annotation accuracy across five methods stands at 0.8807, suggesting that higher CAMUS scores are typically associated with more accurate annotations by the methods. Specifically, for the retina data (**Supplemental Table S1**), 595 out of the 598 data pairs demonstrated a positive correlation between the CAMUS score and the accuracy of the five annotation methods. It showed an average correlation of over 0.74 across all query datasets (**Figure 3a**). Moreover, for all the 672 pairs of cross-species datasets, CAMUS achieved an accuracy of over 0.82 in selecting the best methods (**Figure 3b**). Using methods selected by CAMUS, we observed a median of 51.3% increase in accuracy, a 49.4% increase in macro F1 score, and a 57.3% increase in weighted F1 score compared to the random selection (**Figure 3c**).

**Figure 3.**
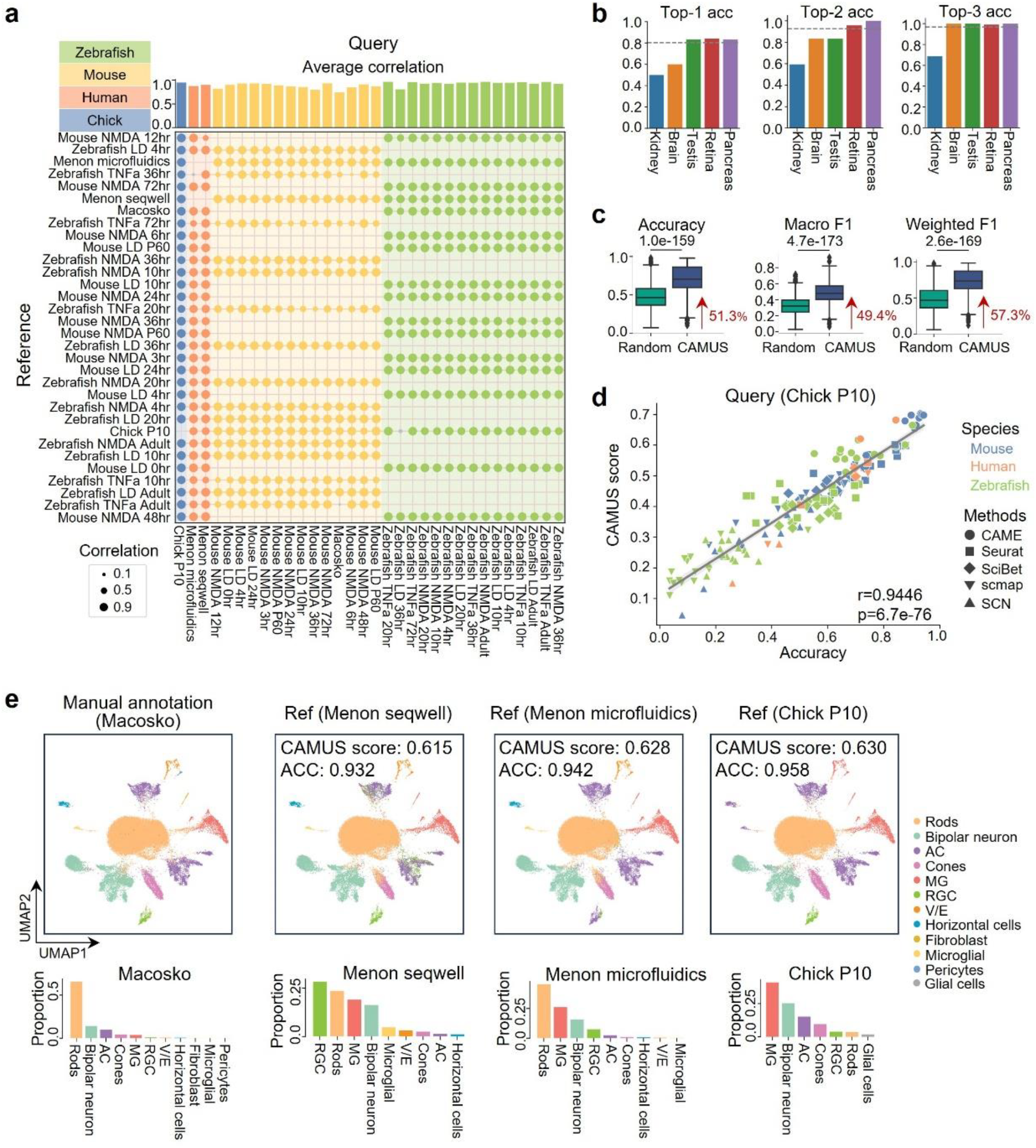
Application of CAMUS for method selection in 672 pairs of cross-species cell type annotation. **a**, Dot plot showing the correlation between the accuracy of five cell annotation methods (i.e., CAME, Seurat, SciBet, scmap, and SCN) and the corresponding CAMUS score for retina data collected from zebrafishes, mice, humans, and chicks. For each dot, color represents a positive correlation, and gray represents a negative correlation. For each dot, color represents a positive correlation, and gray represents a negative correlation. Each cross-species pair has corresponding reference–query combinations, whereas same-species pairs do not. **b**, Bar plot showing the top-1, top-2, and top-3 accuracy of selecting the best methods by CAMUS or randomly. **c**, Boxplot indicating the accuracy, macro F1 score, and weighted F1 score of cell type annotation for methods selected by CAMUS or random. The data following the red arrow represents the percentage improvement for the median value relative to random selection. **d**, Pearson correlation between the CAMUS score and cell type annotation accuracy, where each point represents a reference-method pair. **e**, UMAP plot of mouse retina cells from the macosko dataset, cells are colored according to the manual annotation, and the annotations of CAME using the menon seqwell, menon microfluidics, and chick P10 as references (upper panel). Bar plot showing the cell ratio of the corresponding dataset (lower panel).

Having confirmed that CAMUS can accurately select reference data and methods separately, we next examined the relationship between annotation accuracy and CAMUS scores for all reference-method pairs. For all the 3360 reference-method pairs, we achieved high accuracy in selecting the optimal pair for each query data. Specifically, we attained 49.06% for top-1 accuracy, 63.78% for top-2 accuracy, and 71.70% for top-3 accuracy, respectively. These results represented a significant improvement compared to random selection (**Supplemental Fig. S4a**). Using the Chick P10 dataset (Macosko et al. 2015) as the query and 31 mouse, human, and zebrafish datasets as references (**Figure 3a**), we again observed a strong positive association between the CAMUS score and annotation accuracy across all reference-method pairs (r = 0.9446, p =6.7e-76). Notably, the reference-method pairs that CAMUS ranked highest also produced the most accurate annotations (**Figure 3d**). Meanwhile, in annotating the Macosko dataset using Zebrafish TNFa Adult as the reference, the annotation accuracy of the five methods exhibited a slight negative correlation with the CAMUS scores (**Figure 3a**). Additionally, all methods demonstrated relatively poor annotation accuracy, ranging from 16.42% to 32.37% (**Supplemental Table S2**). To evaluate the cause, we carefully examined the cell proportions and the annotation quality between the reference and query datasets. We identified issues with the annotation of rod cells in the reference data, where some bipolar neurons also expressed the gene *rho*, which is the marker of the rod cell (Hoang et al. 2020) (**Supplemental Fig. S4b, c**). Rod cells happen to be the most prevalent cell type in the Macosko dataset (**Figure 3e**). This indicates that reference quality is a critical factor in the process of reference-based annotation. We found that our clustering of the Macosko dataset resulted in 17 groups, significantly more than the 11 existing cell types (**Figure 3e, Supplemental Fig. S4d**). However, the prioritization of methods based on CAMUS scores still effectively reflected the performance of the annotations. We also discovered that a closer match in cell type proportions between the reference and query data does not necessarily lead to better annotation(**Figure 3e**). Furthermore, the reference data Chick P10, which performed the best with CAME, was not the optimal choice for Seurat (**Supplemental Fig. S4e**). This indicates that different methods have different preferences for the reference data. Therefore, the optimal dataset should be re-selected for each.

### CAMUS achieved additional enhancements when incorporated with reference-integrated strategies

Due to limited reference datasets, we may encounter situations where the annotation may be unsatisfactory even with the best reference-method pair selection. This issue often arises because some references may lack distinctive cell types or be underrepresented, or because some cells are mislabeled. Integrating multiple references should, in principle, widen cell-type coverage and mitigate the impacts of annotation errors in any single dataset. The central question is how to combine multiple references effectively and whether this strategy indeed improves query annotation. To explore this, we collected 3522 pairs of cross-species datasets, where each query data has two distinct references (named reference 1 and 2, respectively) (**STAR Methods**).

We employed the ensemble and multi-reference (multi-ref) strategies to integrate these references. We didn’t apply the ensemble strategy with SciBet and scmap because they do not provide probability-based results (**STAR Methods**). For most methods, employing a multi-ref strategy yields less than a 50% probability of outperforming both single references, i.e., CAME (n=1496, 42.5%), Seurat (n=1384, 39.3%), SCN (n=1333, 37.8%), SciBet (n=1896, 53.8%), and scmap (n=934, 26.5%). We can observe analogous results under an ensemble strategy, i.e., CAME (n=1602, 45.5%), Seurat (n=1499, 42.6%), SCN (n=911, 25.9%) (**Supplemental Fig. S5a**). Moreover, these two strategies ensure that in the majority of instances (∼80% for multi-ref and ∼95% for ensemble), the accuracy will surpass the inferior single reference. In other words, these two strategies enhance annotation accuracy by moderately raising the upper limit and significantly ensuring the lower limit. That’s why multi-ref and ensemble strategies likely yield superior performance compared to randomly selecting a reference (**Supplemental Fig. S5b**). Besides integrating the reference data, another simple approach is to achieve the best-performing reference selection using CAMUS. CAMUS ensures a higher probability that the selected result will surpass any single reference, i.e., CAME (n=2613, 74.2%), Seurat (n=2689, 76.3%), SCN (n=2617, 74.2%), SciBet (n=2550, 72.4%), and scmap (n=2604, 73.9%). CAMUS significantly raises the lower limit of annotation results (**Supplemental Fig. S5a**). We also combined the above three strategies, i.e., using CAMUS to select the best among multi-ref, ensemble, and two single references. We found that this combined approach could yield improvements ranging from 9.3% to 39.3%, compared to randomly selecting from the two individual references (**Supplemental Fig. S5c**).

Next, we annotated the mouse NMDA P60 retinal data (Baron et al. 2016) using the zebrafish LD 10hr retinal data (Baron et al. 2016) (reference 1), zebrafish TNFa 72hr retinal data (Baron et al. 2016) (reference 2), and the two integrating strategies (**Supplemental Fig. S5d**). Using references 1 and 2 individually led to annotation errors for bipolar neurons and rod cells, respectively. The ensemble and multi-ref strategies leveraged the advantages of both reference datasets, thereby correcting the misannotated cell types. Moreover, CAMUS perfectly aligned the ranking of these four scenarios (i.e., reference 1 only, reference 2 only, ensemble, and multi-ref) with their actual annotation performance. Additionally, when the two integrating strategies do not yield improvements, CAMUS can also assist us in rejecting them (**Supplemental Fig. S5e**).

In short, CAMUS can help us determine whether reference integration strategies offer improvements over single references and select the better-performing one (**Supplemental Fig. S5b**). When reference data is limited, CAMUS can incorporate reference-integrated approaches, i.e., the multi-ref and ensemble, to achieve better annotations.

### CAMUS enables highly accurate method selection for scST and scATAC-seq data

We further applied CAMUS to 80 scST and five scATAC-seq datasets as the query data to assess its applicability to other complex scenarios (**STAR Methods**). The scST datasets were derived from brain (Shah et al. 2017; Codeluppi et al. 2018; Moffitt et al. 2018; Eng et al. 2019; Zhang et al. 2021; Zeng et al. 2023; Russell et al. 2024), embryo (Lohoff et al. 2022), retina (Choi et al. 2023), kidney (Liu et al. 2023a), and liver (Liu et al. 2023a) profiled by six distinct spatial technologies (**STAR Methods, Supplemental Table S3**), and the five scATAC-seq datasets were obtained from PBMC (2020), BMMC (Luecken et al. 2021), Cortex (Chen et al. 2019), Skin (Ma et al. 2020), and Mop (Yao et al. 2021). Each query dataset pairs with a well-annotated scRNA-seq dataset as a reference (**STAR Methods, Supplemental Table S4**). The CAMUS score and annotation accuracy in 78 scST datasets showed distinct positive correlations. The two exceptions come from the smFISH embryo 1 and 2 data due to their limited 33 genes (**Supplemental Fig. S6a** and

**Table S3**). CAMUS achieved a median probability of 0.8250 to select the best methods (**Figure 4a**). Even with significant adjustments in clustering resolution (from 0.2 to 0.6), CAMUS consistently maintained robust top-1 accuracy ranging from 82.50% to 86.25% (**Supplemental Fig. S4b**), highlighting its robustness to clustering parameters and underscoring its versatility and reliability in various biological scenarios. It is worth noting that, for the clustering of scST data, the limited number of genes measured may lead to blurry clustering boundaries or different cell types wrongly mixed into a single cluster. Specifically, we focused on the mouse hypothalamic region scST dataset containing 11 cell types (Moffitt et al. 2018). With only 161 profiled genes, the clustering method cannot clearly distinguish excitatory and inhibitory neurons (**Supplemental Fig. S7a, b**). However, this does not affect the efficacy of CAMUS, which still provides method prioritizations that align perfectly with annotation accuracy (**Fig.4b**).

**Figure 4.**
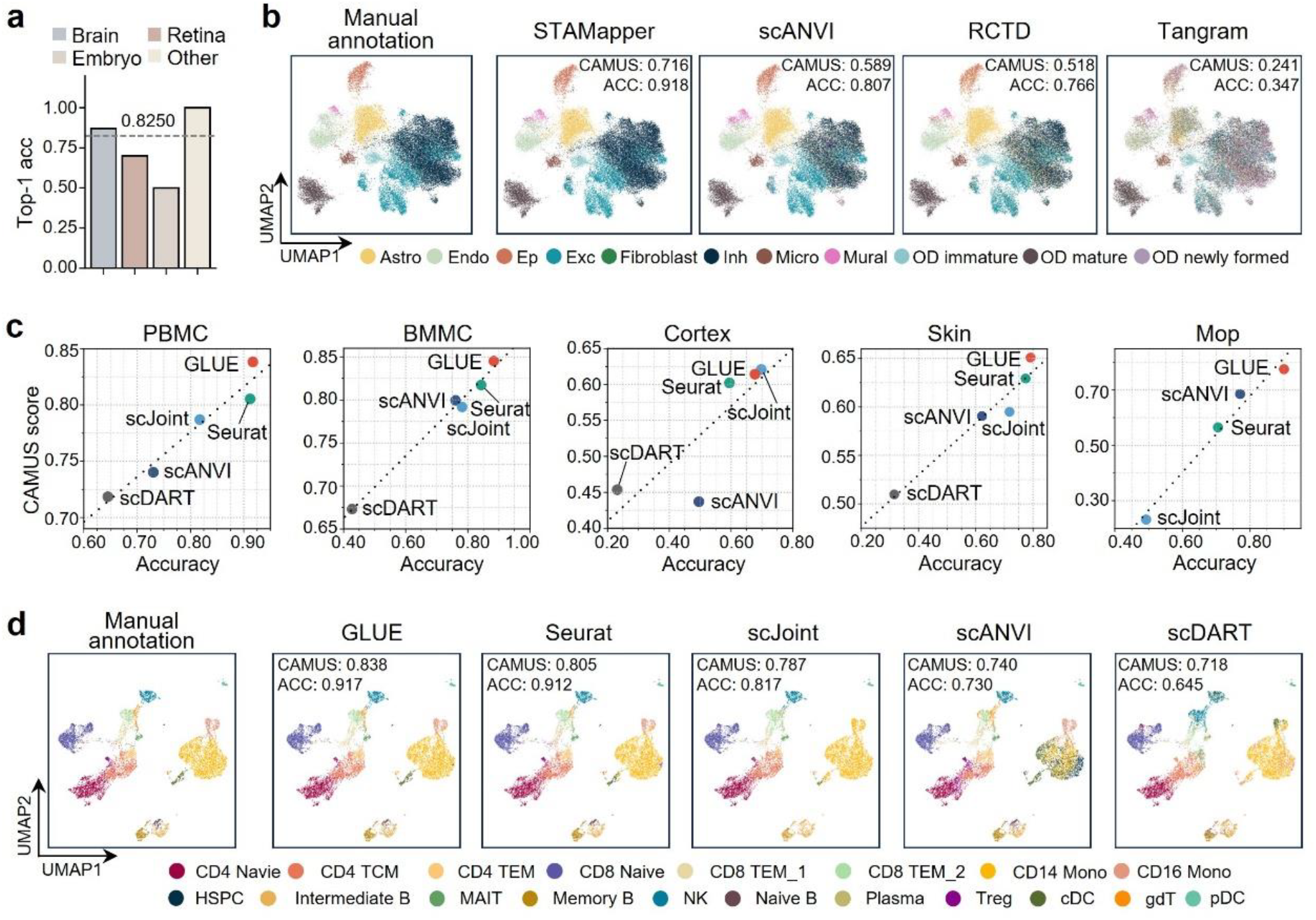
Application of CAMUS is to select annotation methods in cross-modalities and cross-omics cell annotation scenarios. **a**, Bar plot showing the top-1 accuracy of selecting the best methods by CAMUS for annotating scST datasets. **b**, UMAP plots of mouse hypothalamic region scST dataset from the retina, cells are colored by manual annotation, the annotation of STAMapper, scANVI, RCTD, and Tangram, respectively. Astro, astrocytes; Endo, endothelial cells; Ep, ependymal cells; Exc, excitatory neurons; Inh, inhibitory neurons; OD, oligodendrocyte. **c**, Dot plot showing the positive relationship between CAMUS score and ACC. PBMC, Peripheral Blood Mononuclear Cells; BMMC, Bone Marrow Mononuclear Cells; Mop, primary motor cortex. **d**, UMAP plots of PBMC scATAC-seq dataset, cells are colored according to manual annotations, and the annotations by GLUE, Seurat, scJoint, scANVI, and scDART, respectively.

For the scATAC-seq datasets, we applied a higher resolution for the clustering method because their annotations were at the subtype level. Notably, we did not employ scDART for the Mop data due to its runtime exceeding a week (**STAR Methods**). We found that, in most cases, GLUE outperformed other methods, which was consistent with its CAMUS score. scJoint achieved slightly higher accuracy on the cortex data. CAMUS successfully captured this and selected scJoint as the optimal method (**Figure 4c**). CAMUS exhibited perfect top-1 accuracy and stable performance with varied clustering resolution (**Supplemental Fig. S6d**). To further assess CAMUS’s performance at the cellular subtype resolution, we carefully analyzed the PBMC data, which comprises 19 subpopulations. Although CAMUS pre-clustered the data into only 12 clusters, showing a distinct mismatch with the original number of subpopulations, it still achieved perfect consistency with annotation accuracy (**Figure 4d, Supplemental Fig. S6c**). Specifically, GLUE and Seurat exhibited comparable accuracy (0.917 vs. 0.912), while GLUE achieved the higher CAMUS score as expected (0.838 vs. 0.805). In contrast, scJoint mistakenly annotated CD16 Mono cells as CD14 Mono cells, resulting in lower accuracy (0.817) and CAMUS score (0.787). scANVI misclassified a subset of CD8 TEM_2 cells as gdT cells and part of the CD14 Mono cells as cDC and HSPC cells, leading to reduced accuracy (0.730) and CAMUS score (0.740). scDART produced nearly entirely incorrect annotations for CD8 TEM_2 cells, CD16 Mono cells, and gdT cells, yielding the lowest accuracy (0.718) and CAMUS score (0.645).

From a theoretical perspective, both GLUE and Seurat successfully annotated nearly all existing cell types, with discrepancies from the clustering results occurring only in a few rare cell types. Consequently, they achieved the highest CAMUS scores. scJoint, despite misclassifying only one minor subtype (CD16 Mono), still showed a moderate drop in CAMUS score, ranking just behind GLUE and Seurat. On the other hand, scANVI and scDART produced multiple mis-annotations, sharply reducing both accuracy and CAMUS score. For example, scANVI annotated a large fraction of cells in the CD14 Mono population as cDC or HSPC cells, whereas scDART annotated several distinct cell types into NK cells. These errors followed a common pattern: either splitting a single major cell type into several incorrect annotations or merging multiple cell types into one. CAMUS can capture such common-pattern errors in mis-annotations, as they inherently conflict with the underlying clustering structure, thereby assisting researchers in identifying and avoiding them in practical analyses. We also applied CAMUS to four additional scATAC-seq datasets collected from retina, Alzheimer’s disease (AD) cortex, control cortex, and kidney (Morabito et al. 2021; Muto et al. 2021; Wang et al. 2022). CAMUS continues to select the best annotation method, confirming its robustness and generalizability in the scATAC-seq setting (**Supplemental Fig. S7e**).

### CAMUS score helps predict the accuracy of cell type annotation

Although CAMUS facilitates the reference and method selection, users may not know whether the chosen reference-method pair can produce reliable annotations. For instance, if the best reference-method pair achieves a CAMUS score of 0.5, users cannot predict the corresponding annotation accuracy, which may range from ∼20% to ∼90% (**Figure 2a**). When users lack relevant biological knowledge, or when annotating data from a new species, it is exceedingly difficult for them to estimate the annotation accuracy. Even with relevant biological background knowledge, such as verifying the cell type annotations by examining the marker genes’ expression, a significant amount of time is required. Therefore, it becomes critical to provide an estimate of annotation accuracy rapidly.

Given that the CAMUS score significantly correlates with accuracy, it could serve as a powerful predictor. Other predictors, such as the cell number, the cluster number, and cell entropy under different clustering resolutions in the query data, also contribute to predicting the cell annotation accuracy (**STAR Methods**). We adopted the recently published machine learning method AutoGluon (Erickson et al. 2020) and the median absolute error (MAE) between the estimated accuracy and the actual accuracy as the evaluation metric. We achieved an MAE of 0.0453 for the ten-fold cross-validation on 672 pairs of cross-species data encompassing all the cell annotation methods (**Supplemental Fig. S8a, STAR Methods**). Methods that performed better in annotation achieved even lower MAE values: CAME (0.0375), Seurat (0.0448), and SciBet (0.0344) (**Supplemental Fig. S8a, STAR Methods**). Moreover, we validated the trained model on scST data, achieving an MAE of 0.0560 (**Supplemental Fig. S5b**). Our model exhibits low error across all transcriptomic data, potentially attributable to the similarity in their data types and the same normalization procedures. Despite the preprocessing and normalization methods for scATAC-seq data being entirely different from those of scRNA-seq, our model still achieved a low MAE (0.0557) (**Supplemental Fig. S8c**). Consistent with the conclusions drawn from cross-species data, for the best-performing method, GLUE, we achieved a lower MAE (0.0428). This indicated that for the best results obtained through CAMUS selection, we can expect even lower MAEs.

## Discussion

Precise and rapid annotation of single-cell data with reference-based methods is essential for deciphering cell types and their complex interactions (Dimitrov et al. 2022). The selection of references and methods distinctly influences the outcomes of annotations (**Figure 2a, 3c**). However, the critical selection process has not been seriously explored. In this study, we develop a universal reference and method selection method, CAMUS. CAMUS provides a score significantly correlated with annotation accuracy. It works by directly comparing data clustering results with its annotations, thus enabling a smooth expansion to any biologically structured single-cell data.

In real-world datasets, CAMUS demonstrated high accuracy in selecting references and methods for annotating scRNA-seq, scST, and scATAC-seq data. Extensive tests indicate that the clustering resolution has a minimal impact on the performance of CAMUS. Despite unclear boundaries or some errors in clustering, CAMUS consistently achieved high top-1 accuracy. Even when there was a significant discrepancy between the number of clusters and the actual cell types, the performance of CAMUS was not compromised. CAMUS can also be employed for method selection when dealing with annotations of subtypes within the data. In many single-cell datasets, particularly in modalities such as scATAC-seq, ground-truth cell type annotations are either unavailable or uncertain. This limitation poses a challenge for objectively evaluating the reliability of computational annotation methods. Our results demonstrate that CAMUS scores are strongly and positively correlated with annotation accuracy when ground-truth labels are available, suggesting that CAMUS could serve as a proxy metric for annotation quality in settings where such labels are lacking.

The reference data quality is critical for the reference-based annotation of single cells. Moreover, incorrect cell annotations can adversely affect the outcomes of annotation methods. Therefore, users should carefully evaluate the reference data quality and its annotations before employing reference-based methods. CAMUS is a powerful tool for assessing the effectiveness of the reference data for annotating query data. When the reference data is of high quality, the differences among methods diminish significantly. Thus, when sufficient reference datasets are available, users should prioritize selecting the appropriate references. When a specific new single-cell technique lacks mature annotation methods, collecting and selecting proper references through CAMUS can be highly beneficial. This also poses new demands for establishing databases to collect more high-quality datasets (e.g., SpatialDB (Fan et al. 2020) and PanglaoDB (Franzén et al. 2019)).

Recently, an AI-generated reference atlas launched by Synthesize Bio (https://app.synthesize.bio/datasets) has emerged as a promising resource for standardising cell-type annotations across studies. Although the current early-access release is limited to bulk RNA-seq data, the developers have announced that single-cell modalities are in active development. Once available, these synthetic single-cell datasets could provide abundant, cost-effective, bias-reduced cell-type annotations as well as a reproducible reference framework. When integrated with CAMUS, these AI-generated reference atlases should markedly enhance the robustness and cross-study comparability.

## Limitations of the study

One potential limitation of CAMUS is that it does not consider the prior biological knowledge inherent in the data, such as marker genes for its potential cell types, which could serve as an additional measure of annotation accuracy. Moreover, a diverse range of methods and cross-modality datasets should be collected. For instance, transcriptomics to proteomics or transcriptomics to metabolomics, to comprehensively evaluate and improve CAMUS. In the future, a dedicated database of datasets and methods should be established, integrating CAMUS to assist users in selecting reference data and annotation methods.

## STAR Methods

### Data preprocessing

We processed the count matrices for scRNA-seq and scST datasets via Scanpy (Wolf et al. 2018) (https://scanpy.readthedocs.io/en/stable/index.html). We first normalized the library size for each cell, followed by a logarithmic transformation of the expression data with a pseudo-count using the scanpy.pp.normalize_total() function and the scanpy.pp.log1p() function. Subsequently, we selected the top 3000 highly variable genes (HVGs) using the scanpy.pp.highly_variable_genes() function. In cases where the scST datasets comprised fewer than 3000 genes, we used the entire gene set. The data were then scaled using the scanpy.pp.scale() function and subjected to principal component analysis (PCA) via the scanpy.pl.pca() function, where the top 50 principal components were computed. A k-nearest neighbor (KNN) graph was then constructed using 10 neighbors and the first 40 principal components using the scanpy.pp.neighbors() function. Clustering was subsequently performed on the KNN graph using the Leiden algorithm (Traag et al. 2019) with the default resolution parameter set to 0.4 via the scanpy.tl.leiden() function for all datasets presented in **Figure 4d**. We used SnapATAC2 to preprocess the scATAC-seq datasets. Following its documentation (https://kzhang.org/SnapATAC2/index.html), we first imported data fragment files and the reference genome (hg38 for humans and mm10 for mice) using the snap.pp.import_fragments() function. We generated a cell-by-bin matrix containing insertion counts across genome-wide 500-bp bins using the snap.pp.add_tile_matrix() function. We then employed spectral embedding calculated on the top 250k features using the snap.tl.spectral() and snap.pp.select_features() functions. Finally, we clustered the data using the Leiden algorithm with a default resolution parameter of 0.7 and the snap.tl.leiden() function for datasets presented in **Figure 4d**. We also adjusted the clustering resolution for datasets presented in **Supplemental Fig. S7e** (0.1 for the kidney and retina data, 0.3 for the cortex data) (see Section ‘Adjusting the clustering resolution in CAMUS’).

### Benchmarking cell-type annotation

#### Cross-species scRNA-seq datasets

We performed (1) CAME (Liu et al. 2023b) with its Python package, referred to its documentation (https://xingyanliu.github.io/CAME/tut_notebooks/getting_started_pipeline_un.html);

(2) Seurat (Hao et al. 2024) with its R package detailed in its documentation (https://satijalab.org/seurat/articles/integration_mapping); (3) SciBet (Li et al. 2020) with its R package detailed in its documentation (http://scibet.cancer-pku.cn/); (4) scmap-cluster (Kiselev et al. 2018) with the R package scmap, following its documentation (https://bioconductor.org/packages/release/bioc/vignettes/scmap/inst/doc/scmap.html); and (5) SCN (Tan and Cahan 2019) with its R package singleCellNet, following its documentation (https://github.com/CahanLab/singleCellNet?tab=readme-ov-file#introduction).

#### scST datasets

We performed (1) STAMapper with its Python package, referred to its documentation (https://github.com/zhanglabtools/STAMapper/tree/main/Tutorials); (2) scANVI (Xu et al. 2021) with the scVI Python package, referencing the “Integration and label transfer with Tabula Muris” section from its documentation (https://docs.scvi-tools.org/en/stable/tutorials/notebooks/scrna/tabula_muris.html); (3) RCTD (Cable et al. 2022) with the R package spacexr, utilizing its default workflow available at (https://raw.githack.com/dmcable/spacexr/master/vignettes/spatial-transcriptomics.html); and (4) Tangram (Biancalani et al. 2021) with its Python package, employing the cluster mode configuration detailed in the tutorial (https://tangram-sc.readthedocs.io/en/latest/tutorial_sq_link.html).

#### scATAC-seq datasets

We implemented (1) GLUE (Cao and Gao 2022) with its Python package scglue, referring to its documentation (https://scglue.readthedocs.io/en/latest/); (2) scJoint with its Python package detailed in its GitHub page (https://github.com/SydneyBioX/scJoint); (3) Seurat (Hao et al. 2024) with its R package, referring to its documentation (https://satijalab.org/seurat/articles/seurat5_atacseq_integration_vignette); (4) scANVI (Xu et al. 2021), referring to the documentation from SnapATAC2(Zhang et al. 2024) (https://kzhang.org/SnapATAC2/tutorials/annotation.html); (5) scDART (Zhang et al. 2022) with its Python package, following its documentation (https://github.com/PeterZZQ/scDART). Since GLUE and scDART only perform the data integration of scRNA-seq and scATAC-seq data, we trained a k-nearest neighbor (kNN) classifier from the sklearn (Pedregosa et al. 2011) Python package with k=10 on the integrated embeddings of scRNA-seq data to predict the cell labels from scATAC-seq data.

### Calculation of CAMUS score

CAMUS takes the count matrix of query data and the annotation results generated by different references and methods as input. CAMUS consists of two parts, we first pre-cluster the query data (section “Data preprocessing”). For query data with *m* cells: *C* = {*c*_1_, …, *c*_*m*_}, suppose we have the clustering results *U* = {*U*_1_, …, *U*_*s*_} with *s* clusters, where *U*_*i*_ denotes *i*th cluster set and the annotation results *V* = {*V*_1_, …, *V*_*t*_} with *t* cell types, where *V*_*i*_ denotes the cell set of type *i*. It should be noted that for all *i* ≠ *j*:

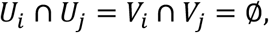

and

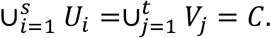

In other words, we can view *U* and *V* as two partitions of *C*.

We then define the concordance score between the two partitions *U* and *V* as the adjusted mutual information (Vinh et al. 2009). Let matrix *M* = [*n*_*ij*_], *i* = 1, …, *s, j* = 1, …, *t*, where *n*_*ij*_ denotes the number of shared cells for *U*_*i*_ and *V*_*j*_. We calculate the probability of randomly picking a cell that belongs to *U*_*i*_:

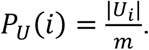

The entropy of the partition *U* is:

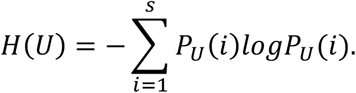

Similarly, we calculate the entropy for partition *V*:

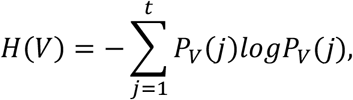

where 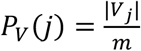. The mutual information between the two partitions is:

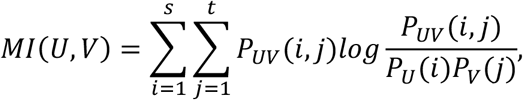

where *P*_*UV*_ (*i, j*) denotes the probability of randomly picking a cell that belongs to *U*_*i*_ ∩ *V*_*j*_, i.e.,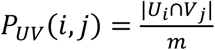. We also calculated the expected mutual information:

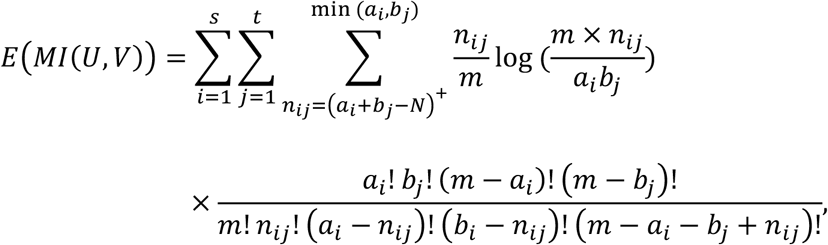

where (*a*_*i*_ + *b*_*j*_ − *N*) denotes max (0, *a*_*i*_ + *b*_*j*_ − *N*) and 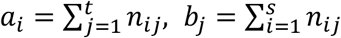.

Finaly, we defined the CAMUS concordance score between *U* and *V* as:

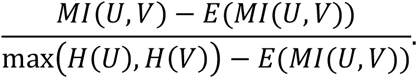

To improve consistency and align with common practice, we use the arithmetic mean of *H*(*U*), *H*(*V*) instead of the max. The score is assigned a value of one when the two partitions are identical, and it takes a value of zero when the mutual information between the two partitions equals the value expected by chance alone (**Supplemental Note 1**). The CAMUS score approaches one as the alignment between the pre-cluster and the annotation results improves. Conversely, the CAMUS score tends toward zero when there is a marked discrepancy between these results.

### Adjusting the clustering resolution in CAMUS

Conceptually, the performance of CAMUS is most reliable when the number of clusters is close to the number of annotated cell types, as the CAMUS score is fundamentally derived from the concordance between clustering results and annotations. To achieve this, we recommend adjusting the resolution parameter in CAMUS to align the granularity of the annotation level derived from reference-based methods. The resolution parameter influences the tendency of the pre-clustering step to generate finer or coarser partitions: higher values encourage the formation of smaller, more detailed clusters, whereas lower values favor larger, more aggregated clusters.

### Estimate the annotation accuracy

We used the number of total cells of the query dataset as the first predictor. We also used the number of clusters, cell entropy, and CAMUS concordance score under the clustering resolution of 0.2, 0.4, 0.6, and 0.8 as other predictors. Suppose we have a dataset of *m* cells, and clustering results *U* = {*U*_1_, …, *U*_*s*_}, where the number of cells in *U*_*i*_ is *m*_*i*_. To calculate the cell entropy, we first calculate the cell ratio for each cluster: 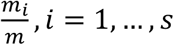, *i* = 1, …, *s*, and then the cell entropy as:

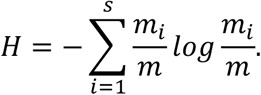

We applied AutoGuon to 13 predictors to estimate the annotation accuracy, we set eval_metric=‘median_absolute_error’ for the “TabularPredictor” function, and set time_limit=1800, presets=‘best_quality’ for the “fit” function.

### Reference-integrated strategies

Suppose we have a query dataset with *m* cells and two reference datasets: reference 1 with *n*_1_ cells and *p*_1_ genes, and reference 2 with *n*_2_ cells and *p*_2_ genes. Here, we propose two strategies to integrate these two references. For the references of the same species and tissue type, we extract all possible combinations of them, ultimately obtaining 3,522 data triplets (reference 1, reference 2, query).

#### Ensemble

Suppose we have *q*_1_ distinct cell types for reference 1 and *q*_2_ distinct cell types for reference 2. Both references share *q* common cell types. Then, using reference 1 and query datasets as the input of the cell annotation method, we will have a probability matrix 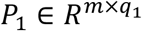, where each the row *i* and column *j* of *P*_1_ denotes the probability of the cell *i* belongs to cell type *j*. And we will have 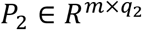 when using reference 2 and query datasets as the input. We denote 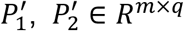 as the probability matrix for the *q* shared cell types. Also, utilizing the annotation results from references 1 and 2, we can calculate the CAMUS scores *u*_1_ and *u*_2_, respectively. We then ensemble the two probability matrices:

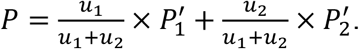

We next incorporate the unique cell type probabilities to matrix *P* and assign the cell type with the highest probability to the corresponding cells.

#### Multi-ref

Here, we concatenate the two reference datasets. Assuming that reference 1 and 2 datasets share *p* genes, the resulting combined matrix consists of *n*_1_ + *n*_2_ cells and *p* genes. We used the combined matrix as a new reference to annotate query data.

## Data availability

All datasets analyzed in this study are publicly available from the links provided.

### Cross-species scRNA-seq datasets

Dataset 1-2: (Pancreas, Human, Mouse), inDrop, GSE84133 in the GEO database.

Dataset 3: (Pancreas, Human), CelSeq, CelSeq2, Fluidigm C1, and SMART-Seq2, a mixed pancreas dataset produced across four technologies, distributed in SeuratData package, https://satijalab.org/seurat/articles/integration_mapping.

Dataset 4: (Pancreas, Mouse), Smart-seq2, https://figshare.com/articles/dataset/Single-cell_RNA-seq_data_from_Smart-seq2_sequencing_of_FACS_sorted_cells_v2_/5829687/7.

Dataset 5-7: (Testis, Human, Monkey, Mouse), Drop-seq, GSE142585 in the GEO database.

Dataset 8: (Brain, Human), snDrop-seq, GSE97942 in the GEO database.

Dataset 9: (Brain, Mouse), Smart-seq2, GSE115746 in the GEO database.

Dataset 10: (Brain, Mouse), Drop-seq, GSE93374 in the GEO database.

Dataset 11: (Brain, Mouse), Smart-seq2, https://figshare.com/articles/dataset/Single-cell_RNA-seq_data_from_Smart-seq2_sequencing_of_FACS_sorted_cells_v2_/5829687/7.

Dataset 12-13: (Brain, Lizard, Turtle), Drop-seq, https://brain.mpg.de/research/laurent-department/software-techniques.html.

Dataset 14: (Kidney, Mouse), Drop-seq, GSE94333 in the GEO database.

Dataset 15: (Kidney, Mouse), Drop-seq, GSE111107 in the GEO database.

Dataset 16: (Kidney, Mouse), 10x, GSE107585 in the GEO database.

Dataset 17: (Kidney, Mouse), 10x, https://figshare.com/articles/dataset/Single-cell_RNA-seq_data_from_microfluidic_emulsion_v2_/5968960/2.

Dataset 18: (Kidney, Human), 10x snRNA-seq, GSE118184 in the GEO database.

Dataset 19: (Kidney, Human), 10x, EGAS00001002171, EGAS00001002486, EGAS00001002325, and EGAS00001002553 in European Genome-phenome Archive.

Dataset 20: (Kidney, Human), 10x, GSE114530 in the GEO database.

Dataset 21: (Kidney, Human), 10x, GSE109488 in the GEO database.

Dataset 22: (Retina, Human), microfluidics, GSE137847 in the GEO database.

Dataset 23: (Retina, Human), Seq-Well, GSE137847 in the GEO database.

Dataset 24: (Retina, Mouse), Drop-seq, GSE63473 in the GEO database.

Dataset 25-37: (Retina, Mouse), 10x, GSE135406 in the GEO database.

Dataset 38-53: (Retina, Zebrafish), 10x, GSE13540 in the GEO database.

Dataset 54: (Retina, Chick), 10x, GSE13540 in the GEO database.

A detailed summary of the 672 reference-query pairs can be found in **Table S1**.

### Paris of scRNA-seq and scST datasets

We collected 80 pairs of scRNA-seq and scST datasets from https://github.com/zhanglabtools/STAMapper/. A human liver cancer dataset profiled by NanoString was excluded due to the absence of manual annotation.

### Paris of scRNA-seq and scATAC-seq datasets

Dataset 1 (PBMC, Human), 10x Multiome, https://support.10xgenomics.com/single-cell-multiome-atac-gex/datasets/1.0.0/pbmc_granulocyte_sorted_10k?.

Dataset 2 (BMMC, Human), 10x Multiome, GSE194122 in the GEO database.

Dataset 3 (Cortex, Mouse), SHARE-seq, GSE126074 in the GEO database.

Dataset 4 (Skin, Mouse), SHARE-seq, GSE140203 in the GEO database.

Dataset 5 (Mop, Mouse), The 10x scRNA-seq and scATAC-seq datasets can be accessed from NeMO archive https://assets.nemoarchive.org/dat-ch1nqb7. We used samples with id: CEMBA171206_3C, CEMBA171207_3C, CEMBA171213_4B, CEMBA180104_4B, CEMBA180409_2C, CEMBA180410_2C, CEMBA180612_5D, CEMBA180618_5D.

Dataset 6 (Retina, Human), 10x Multiome, GSE196235 in the GEO database.

Dataset 7, 8 (Normal and Alzheimer’s Disease Cortex, Human), 10x scRNA-seq and single-nucleus ATAC-seq (snATAC-seq). GSE174367 in the GEO database.

Dataset 9 (Kidney, Human), 10x snRNA-seq and scATAC-seq. GSE151302 in the GEO database.

## Code availability

CAMUS is implemented in Python and is available at GitHub [https://github.com/zhanglabtools/CAMUS] and as Supplemental Code.

## Acknowledgments

This work has been supported by the National Key Research and Development Program of China (no. 2021YFC2701601 to S.Q.Z.), the Science and Technology Commission of Shanghai Municipality (no. 23JC1401000 to S.Q.Z.), the National Natural Science Foundation of China (nos. 32341013, 12326614 to S.Z., and 12471350 to S.Q.Z.), the R&D project of Pazhou Lab (Huangpu) (no. 2023K0602 to S.Z.) and the CAS Project for Young Scientists in Basic Research (no. YSBR-034 to S.Z.).

## Notes

### Competing Interest Statement

The authors have declared no competing interest.

### Summary of Updates

We change the title and more detailed test.

