## Supplemental Text and Figures for "Highly accurate reference and method selection for universal cross-dataset cell type annotation with CAMUS"

**Supplemental Note 1** We used the clustering results $U=\{U_1,\dots,U_s\}$ with $s$ clusters and the annotation results $V=\{V_1,\dots,V_t\}$ with $t$ cell types to calculate the CAMUS concordance score formulated as follows:

$$\frac{MI(U,V)-E(MI(U,V))}{\max\big(H(U),H(V)\big)-E(MI(U,V))},$$

where

$$H(U)=-\sum_{i=1}^{s}P_U(i)logP_U(i).$$

$$H(V)=-\sum_{j=1}^{t}P_V(j)logP_V(j).$$

Here $P_U(i)=\frac{|U_i|}{m}$, $P_V(j)=\frac{|V_j|}{m}$ and $U_i\cap U_j=V_i\cap V_j=\emptyset$.

$$MI(U,V)=\sum_{i=1}^{s}\sum_{j=1}^{t}P_{UV}(i,j)log\frac{P_{UV}(i,j)}{P_U(i)P_V(j)},$$

where $P_{UV}(i,j)=\frac{|U_i\cap V_j|}{m}$.

$$E\big(MI(U,V)\big)=\sum_{i=1}^{s}\sum_{j=1}^{t}\sum_{n_{ij}=(a_i+b_j-N)^+}^{\min(a_i,b_j)}\frac{n_{ij}}{m}\log\left(\frac{m\times n_{ij}}{a_ib_j}\right)$$

$$\times\frac{a_i!\,b_j!\,(m-a_i)!\,(m-b_j)!}{m!\,n_{ij}!\,(a_i-n_{ij})!\,(b_i-n_{ij})!\,(m-a_i-b_j+n_{ij})!},$$

where $(a_i+b_j-N)^+$ denotes $\max(0,a_i+b_j-N)$ and $a_i=\sum_{j=1}^{t}n_{ij}$, $b_j=\sum_{i=1}^{s}n_{ij}$.

We next prove that the score is assigned a value of 1 when the two partitions are identical, and it takes a value of 0 when the mutual information between the two partitions is equal to the value expected by chance alone. The score is greater than 0 and less than 1.

When $U$, $V$ are two identical partitions, for any $i$, we have:

$$P_U(i)=\frac{|U_i|}{m}=P_V(i)=\frac{|V_i|}{m},$$

when $i\neq j$, we have:

$$P_{UV}(i,j)=\frac{|U_i\cap V_j|}{m}=0,$$

since $U_i\cap V_j=U_i\cap U_j=\emptyset$. When $i=j$, we have:

$$P_{UV}(i,i)=\frac{|U_i\cap V_i|}{m}=\frac{|U_i\cap U_i|}{m}=\frac{|U_i|}{m}=P_U(i)=\frac{|V_i|}{m}=P_V(i),$$

then

$$H(U) = -\sum_{i=1}^{s} P_U(i) log P_U(i) = -\sum_{j=1}^{t} P_V(j) log P_V(j) = H(V).$$

Then the mutual information between the two partitions is:

$$MI(U,V) = \sum_{i=1}^{s}\sum_{j=1}^{t} P_{UV}(i,j) log \frac{P_{UV}(i,j)}{P_U(i)P_V(j)} = \sum_{i=1}^{s} P_{UV}(i,i) log \frac{P_{UV}(i,i)}{P_U(i)P_V(i)}$$

$$= \sum_{i=1}^{s} P_{UV}(i,i) log \frac{P_{UV}(i,i)}{P_U(i)P_V(i)} = \sum_{i=1}^{s} P_U(i) log \frac{P_U(i)}{P_U(i)P_U(i)} = \sum_{i=1}^{s} P_U(i) log \frac{1}{P_U(i)}$$

$$= -\sum_{i=1}^{s} P_U(i) log P_U(i) = H(U).$$

The concordance score is formulated as:

$$\frac{MI(U,V) - E(MI(U,V))}{\max(H(U),H(V)) - E(MI(U,V))} = \frac{H(U) - E(MI(U,V))}{H(U) - E(MI(U,V))} = 1.$$

That is to say, the score is assigned a value of 1 when the two partitions are identical. $E(MI(U,V)$ represents the expected value of $MI(U,V)$ under completely random partitions, averaged over all possible partition assignments that respect the marginal distributions of the data. When the mutual Information between the two partitions is equal to the value expected by random chance. In other words, when $MI(U,V) = E(MI(U,V))$, the concordance score is 0.

To prove the non-negative property of the concordance score, we first prove the non-negative property of $MI(U,V)$. Since we have:

$$\sum_{i=1}^{s}\sum_{j=1}^{t} P_{UV}(i,j) = 1,$$

and the negative logarithm is convex. Therefore, by applying Jensen inequality, we will have:

$$MI(U,V) = \sum_{i=1}^{s}\sum_{j=1}^{t} P_{UV}(i,j) log \frac{P_{UV}(i,j)}{P_U(i)P_V(j)} = -\sum_{i=1}^{s}\sum_{j=1}^{t} P_{UV}(i,j) log \frac{P_U(i)P_V(j)}{P_{UV}(i,j)}$$

$$\geq -\sum_{i=1}^{s}\sum_{j=1}^{t} log P_{UV}(i,j) \times \frac{P_U(i)P_V(j)}{P_{UV}(i,j)} = -\, log \sum_{i=1}^{s}\sum_{j=1}^{t} P_U(i)P_V(j)$$

$$= -\log\left(\sum_{i=1}^{s} P_U(i) \sum_{j=1}^{t} P_V(j)\right) = 0.$$

Since $E(MI(U,V))$ is the expectation of $MI(U,V) \geq 0$, so we will have $E\big(MI(U,V)\big) \geq 0$. Since $\max\big(H(U),H(V)\big)$ is a upper bound for $MI(U,V)$(Ash 2012), we will have:

$$\max\big(H(U), H(V)\big) \geq MI(U,V).$$

Since $E(MI(U,V))$ represents the expected value of mutual information under randomized conditions. We will have:

$$MI(U,V) \geq E(MI(U,V)).$$

That is to say:

$$\max\big(H(U), H(V)\big) \geq E(MI(U,V)).$$

Thus, we have:

$$\frac{MI(U,V) - E(MI(U,V))}{\max\big(H(U), H(V)\big) - E(MI(U,V))} \geq 0.$$

**Supplemental Note 2**

We next compared AMI with Adjusted Rand Index (ARI) and Fowlkes Mallows Index (FMI). We aimed to demonstrate that among these three metrics, AMI is the most suitable for our use cases, as it remains the most robust when there is a substantial discrepancy between the number of pre-clusters and the number of annotated cell types.

The ARI ranges from -1 to 1, while AMI and FMI both range from 0 to 1. To analyze these three indicators at a theoretical level, we designed the following scenario.

For total $N = kn$ samples, suppose we have a pre-clustering result:

$$U = \{U_1, U_2, \dots, U_k\},$$

with $|U_i| = n$ for every $i \in \{1, . , , , k\}$ . Also, we have another clustering/annotation result:

$$V = \{U_{11}, \dots, U_{1t}, U_{21}, \dots, U_{2t}, \dots, U_{k1}, \dots, U_{kt}\} = \{V_1, \dots, V_{kt}\}.$$

Here, we evenly split each $U_i$ into $t$ sub-clusters. Therefore, we have $kt$ clusters, each with $n/t$ samples. Here, we assume that $t$ is a factor of $n$, which means $n/t$ is an integer.

Let matrix $M = \left[n_{ij}\right],\ i = 1, \dots, s,\ j = 1, \dots, t$ , where $n_{ij}$ denotes the number of shared cells for $U_i$ and $V_j$. We first compute the mutual information $MI(U,V)$:

$$P(X \in U_i \cap V_j) = p_{UV}(i,j) = \frac{n_{ij}}{kn} = \frac{n/t}{kn} = \frac{1}{kt},$$

$$P(X \in U_i) = p_U(i) = \frac{n}{kn} = \frac{1}{k},$$

$$P(X \in V_i) = p_V(i) = \frac{1}{kt}.$$

$$MI(U,V) = \sum_{i=1}^{k}\sum_{j=1}^{t} P_{UV}(i,j) log \frac{P_{UV}(i,j)}{P_U(i)P_V(j)} = \sum_{i=1}^{k}\sum_{j=1}^{t} p_{ij} log \frac{p_{ij}}{p_i p_j}$$

$$= kt * \frac{1}{kt} * log\frac{1/kt}{(1/k)*(1/kt)} = logk.$$

$$H(U) = -\sum_{i=1}^{k} P_U(i)logP_U(i) = -k * \frac{1}{k} * \log\frac{1}{k} = logk.$$

$$H(V) = -\sum_{i=1}^{kt} P_v(i)logP_v(i)$$

$$= -kt * \frac{1}{kt} log\frac{1}{kt} = logkt = logk + logt = H(U) + logt.$$

$E\big(MI(U,V)\big)$ is a constant, suppose it equals to $c$.

$$AMI = \frac{MI(U,V) - E(MI(U,V))}{0.5\big(H(U)+H(V)\big) - E(MI(U,V))} = \frac{2logk - c}{2logk + logt - c}$$

$$= \frac{2 - \frac{c}{logk}}{2 + \frac{logt}{logk} - \frac{c}{logk}}.$$

The ARI is formulated as:

$$\mathrm{ARI} = \frac{\sum_{i,j}\binom{n_{ij}}{2} - \frac{\left(\sum_i\binom{a_i}{2}\right)\left(\sum_j\binom{b_j}{2}\right)}{\binom{N}{2}}}{\frac{1}{2}\left[\sum_i\binom{a_i}{2} + \sum_j\binom{b_j}{2}\right] - \frac{\left(\sum_i\binom{a_i}{2}\right)\left(\sum_j\binom{b_j}{2}\right)}{\binom{N}{2}}}.$$

Here, $n_{ij} = |U_i \cap V_j|$, $a_i = \sum_j n_{ij}$, $b_j = \sum_i n_{ij}$, $N = \sum_{ij} n_{ij}$. Utilize the same definition of $U$ and $V$ in AMI. We have:

$$\sum_{ij}\binom{n_{ij}}{2} = kt\binom{n/t}{2} = \frac{k}{2}\left(\frac{n^2}{t} - n\right),$$

$$\sum_i \binom{a_i}{2} = k\binom{n}{2} = \frac{k}{2}(n^2 - n), \quad \sum_j \binom{b_j}{2} = kt\binom{\frac{n}{t}}{2} = \frac{k}{2}\left(\frac{n^2}{t} - n\right),$$

$$\binom{N}{2} = \binom{kn}{2} = \frac{1}{2}(k^2n^2 - kn),$$

$$E = \frac{\left(\sum_i \binom{a_i}{2}\right)\left(\sum_j \binom{b_j}{2}\right)}{\binom{N}{2}} = \frac{1}{2}\cdot\frac{\frac{n^4}{t} - n^3\left(1 + \frac{1}{t}\right) + n^2}{n^2 - \frac{n}{k}}.$$

When $n$ is sufficiently large, we have:

$$\sum_{ij}\binom{n_{ij}}{2} - E \approx \frac{(k-1)n^2}{2t},$$

$$\frac{1}{2}\left[\sum_{i}\binom{a_i}{2}+\sum_{j}\binom{b_j}{2}\right]-E\approx n^2\left(\frac{k}{4}+\frac{1}{t}\left(\frac{k}{4}-\frac{1}{2}\right)\right).$$

We have:

$$ARI\approx\frac{2(k-1)}{kt+(k-2)}.$$

When $t$ is sufficiently large,

$$ARI\approx\frac{2(k-1)}{k}\frac{1}{t}.$$

The Fowlkes-Mallows score is defined as:

$$FMI=\frac{TP}{\sqrt{(TP+FP)(TP+FN)}}.$$

$TP$: number of pairs that are in the same cluster in both $U$ and $V$.
$FP$: number of pairs that are in the same cluster in $V$ but different clusters in $U$.
$FN$: number of pairs that are in the same cluster in U but different clusters in $V$.
Since each $V_{i,j}\subseteq U_i$, we have:

$$FP=0.$$

That is to say:

$$FMI=\sqrt{\frac{TP}{TP+FN}}.$$

Pairs counted as $TP$ come from the interiors of all $V_{ij}$:

$$TP=\sum_{i=1}^{k}\sum_{j=1}^{t}\binom{n/t}{2}=kt\binom{n/t}{2}.$$

Pairs that are same in $U$ total $k\binom{n}{2}$. Those already in $TP$ are within $V_{ij}$. So

$$FN=k\binom{n}{2}-kt\binom{n/t}{2}.$$

We have:

$$\frac{\mathrm{TP}}{\mathrm{TP}+\mathrm{FN}}=\frac{kt\binom{n/t}{2}}{k\binom{n}{2}}=\frac{t\binom{n/t}{2}}{\binom{n}{2}}=\frac{t\frac{n(n-t)}{2t^2}}{\frac{n(n-1)}{2}}=\frac{n-t}{t(n-1)}.$$

Hence, we have:

$$FMI=\sqrt{\frac{n-t}{t(n-1)}}=\frac{1}{\sqrt{t}}\sqrt{\frac{n-t}{n-1}}=\sqrt{\frac{1-\frac{t}{n}}{1-\frac{1}{n}}}.\frac{1}{\sqrt{t}}.$$

When $n\to\infty, t\to\infty$, and $t=o(n)$, we have:

$$\sqrt{\frac{1-\frac{t}{n}}{1-\frac{1}{n}}} \longrightarrow 1.$$

$$\sqrt{\frac{1-\frac{t}{n}}{1-\frac{1}{n}}}.\frac{1}{\sqrt{t}} \longrightarrow \frac{1}{\sqrt{t}}.$$

In conclusion:

$$AMI = \frac{2-\frac{c}{logk}}{2+\frac{logt}{logk}-\frac{c}{logk}},$$

when $n \to \infty$, we have:

$$\mathrm{ARI} \approx \frac{2(k-1)}{k}\frac{1}{t},$$

when $n \to \infty, t \to \infty,$ and $t = o(n)$:

$$\mathrm{FMI} = \sqrt{\frac{1-\frac{t}{n}}{1-\frac{1}{n}}}.\frac{1}{\sqrt{t}} \longrightarrow \frac{1}{\sqrt{t}}.$$

We found that obtaining an approximate value for the AMI requires minimal conditions, with the only requirement being a sufficiently large total number of single cells. Moreover, when $t$ increases, AMI decays the slowest, FMI next, and ARI is the fastest.

We verified this in a simulated study, where we construct a synthetic dataset comprising three perfectly balanced ground-truth clusters of equal size (500 observations each; 1,500 in total). Predicted partitions are generated by refining each ground-truth cluster into an increasing number of equal-sized subclusters. Specifically, we divided the ground-truth clusters into 1, 2, 5, and 10 parts, yielding predicted cluster counts of 3, 6, 15, and 30, respectively. Importantly, this procedure introduces no cross-cluster mixing: each predicted subcluster is a strict subset of a single true cluster.

For each refinement level, we compare the predicted labels with the ground truth using ARI, AMI, and FMI. We observe that ARI and FMI typically exhibit a steeper decline than AMI (**Supplemental Table S5**), which corroborates our theoretical analysis.

### Supplemental Figures

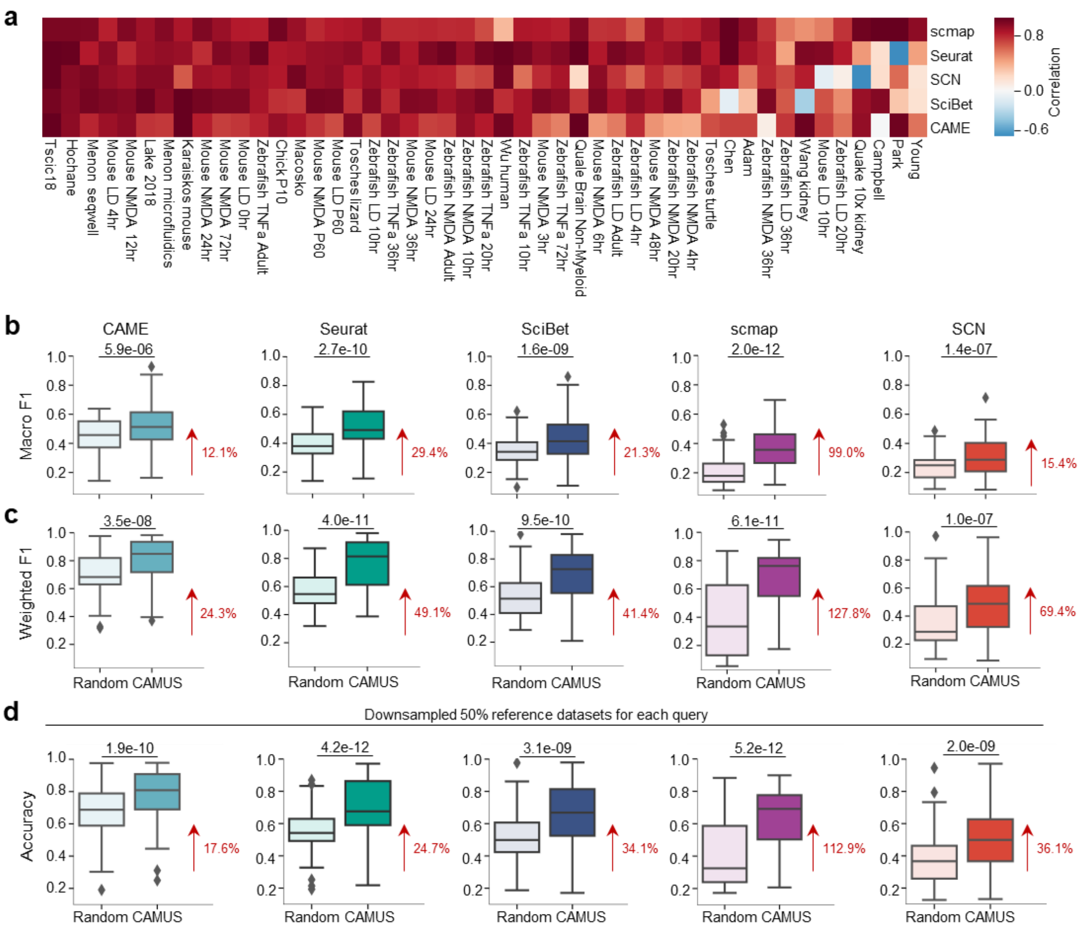


**Supplemental Figure S1. a**, Heatmap showing the correlation of annotation accuracy using different references and corresponding CAMUS scores across five cell-type annotation methods (row) for 47 query datasets (column). Only query datasets with ≥3 references are included. **b-c**, Boxplots comparing the accuracy of annotation achieved by reference random selection versus CAMUS-guided selection under each integration strategy. Percent median improvements over random selection are annotated with red arrows. For the random selection, we performed five independent replicates for each setting and report the average values. The p-value was calculated using the paired t-test. **d**, Boxplot indicating the Accuracy of cell type annotations, where references are selected by CAMUS or randomly. We here. For the random selection, we performed five independent replicates for each setting and report the average values. For each query dataset, we randomly selected 50% of the reference data for annotation. This procedure was repeated independently five times, and the average values were reported. The p-value was calculated using the paired t-test. The data following the red arrow represents the percentage improvement for the median value relative to random selection.

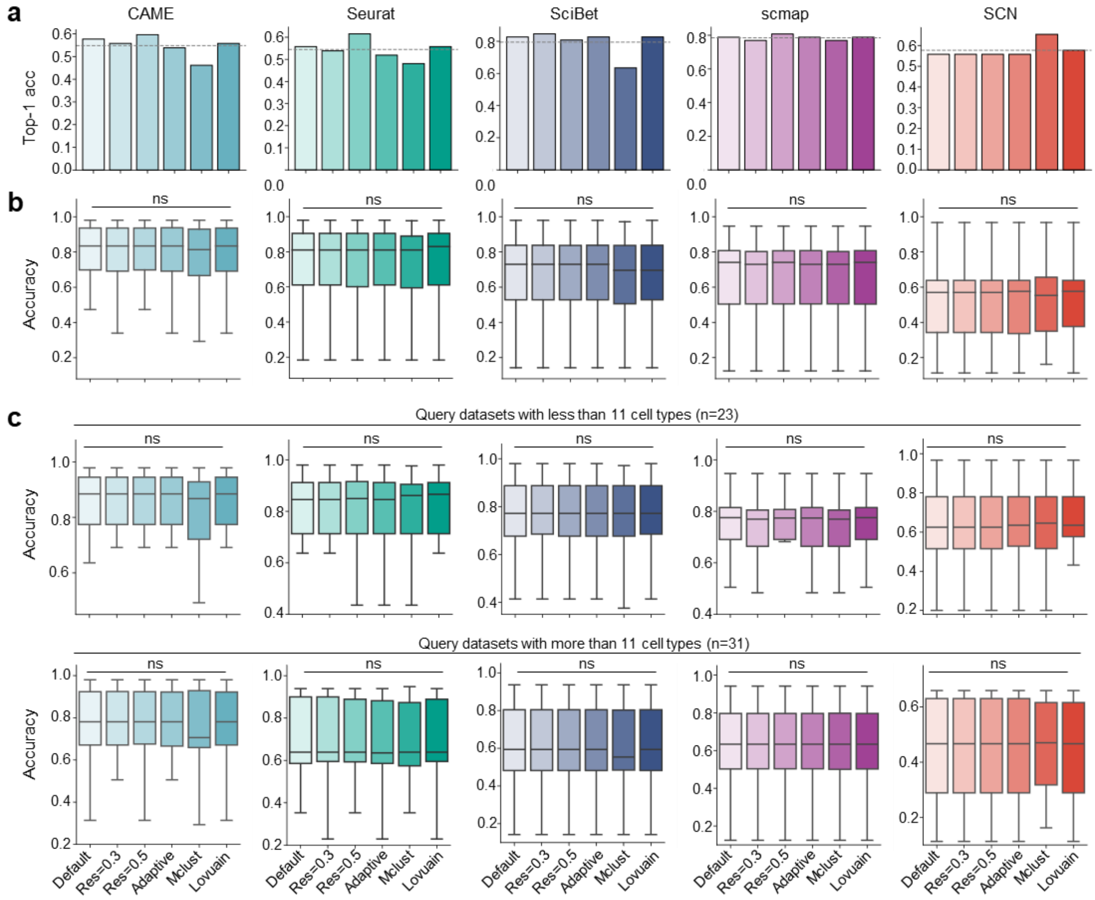


**Supplemental Figure S2. a**, Bar plot showing the top-1 accuracy of selecting the best reference by CAMUS. The gray dashed line indicates the average across diverse settings. **b**, Boxplot showing the annotation accuracy differences using CAMUS to select references across diverse settings. “Adaptive” refers to automatically selecting, from the range of resolution values from 0.10 to 0.95 in increments of 0.05, the one that yields the highest silhouette score. P values were calculated by one-way ANOVA. **c**. We divided the query datasets into those with fewer than 11 cell types (upper panel) and those with more than 11 cell types (lower panel), while keeping the settings for the other settings consistent with those in **(b)**.

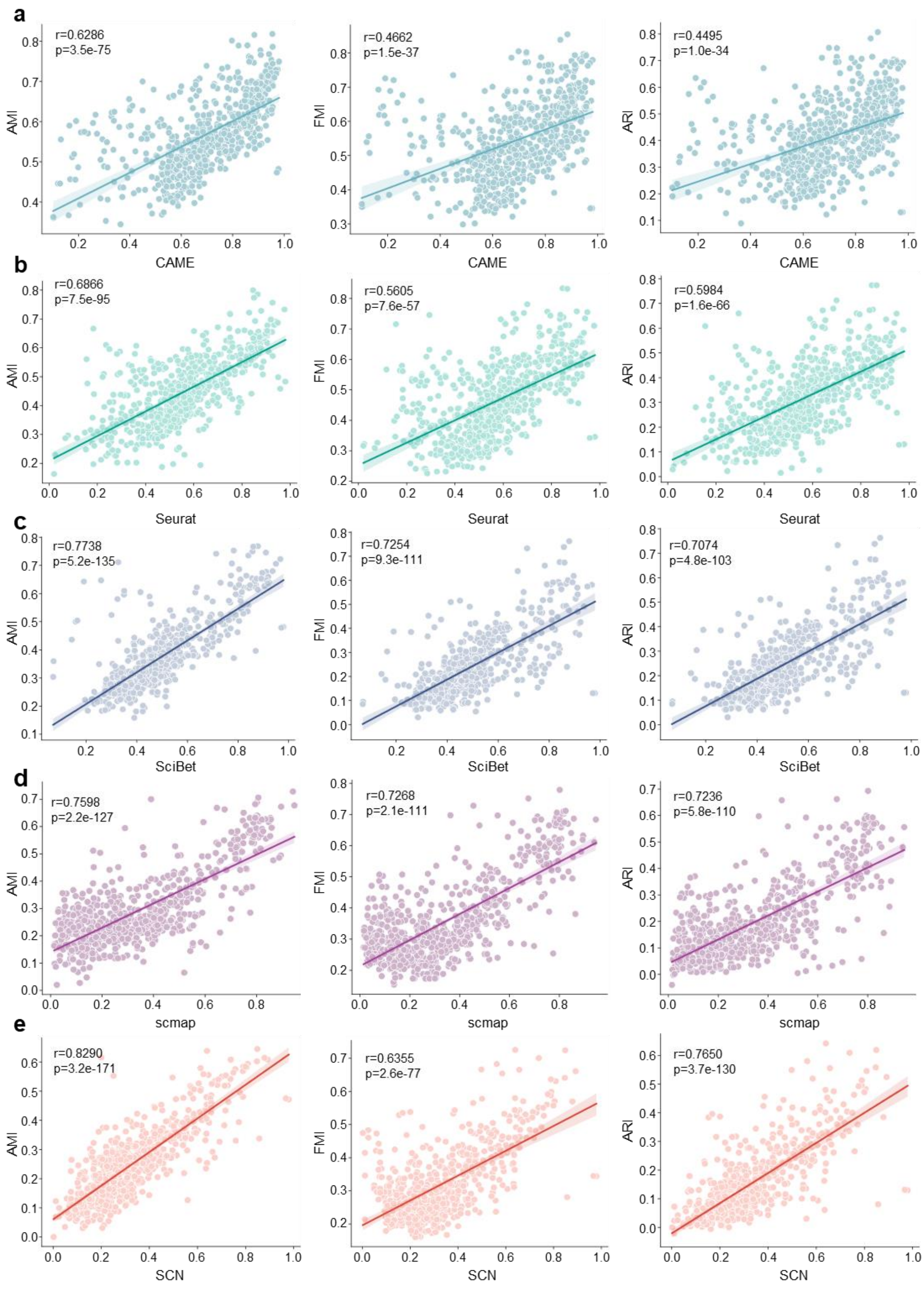


**Supplemental Fig S3.** Scatter plot showing the Pearson's correlation between the AMI, FMI, ARI score, and annotation accuracy in all cross-species data pairs (n=672).

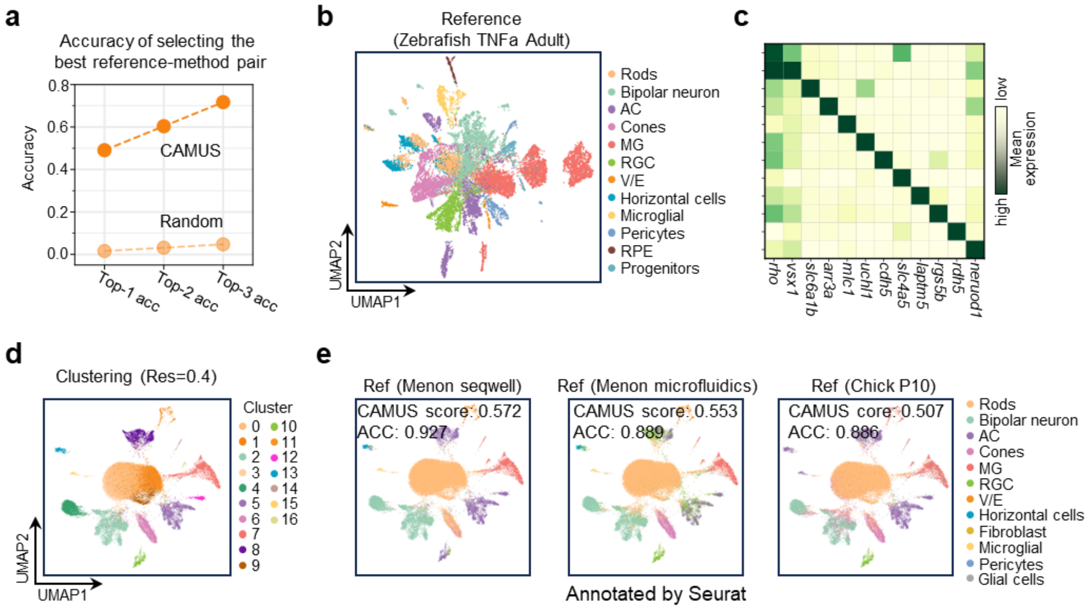


**Supplemental Figure S4. a,** Dot plot showing the top-1, top-2, and top-3 accuracy of selecting the best reference-method pair by CAMUS or randomly. **b**, UMAP plot of zebrafish TNFa adult dataset from the retina, cells are colored by manual annotation. **c,** Heatmap showing the marker gene expression for cell types from the zebrafish TNFa adult dataset. **d,** UMAP plot of mouse retina cells from the macosko dataset, cells are colored by the clustering results. **e,** UMAP plot of mouse retina cells from the macosko dataset, cells are colored according to the annotation result of Seurat using the menon seqwell, menon microfluidics, and chick P10 as references.

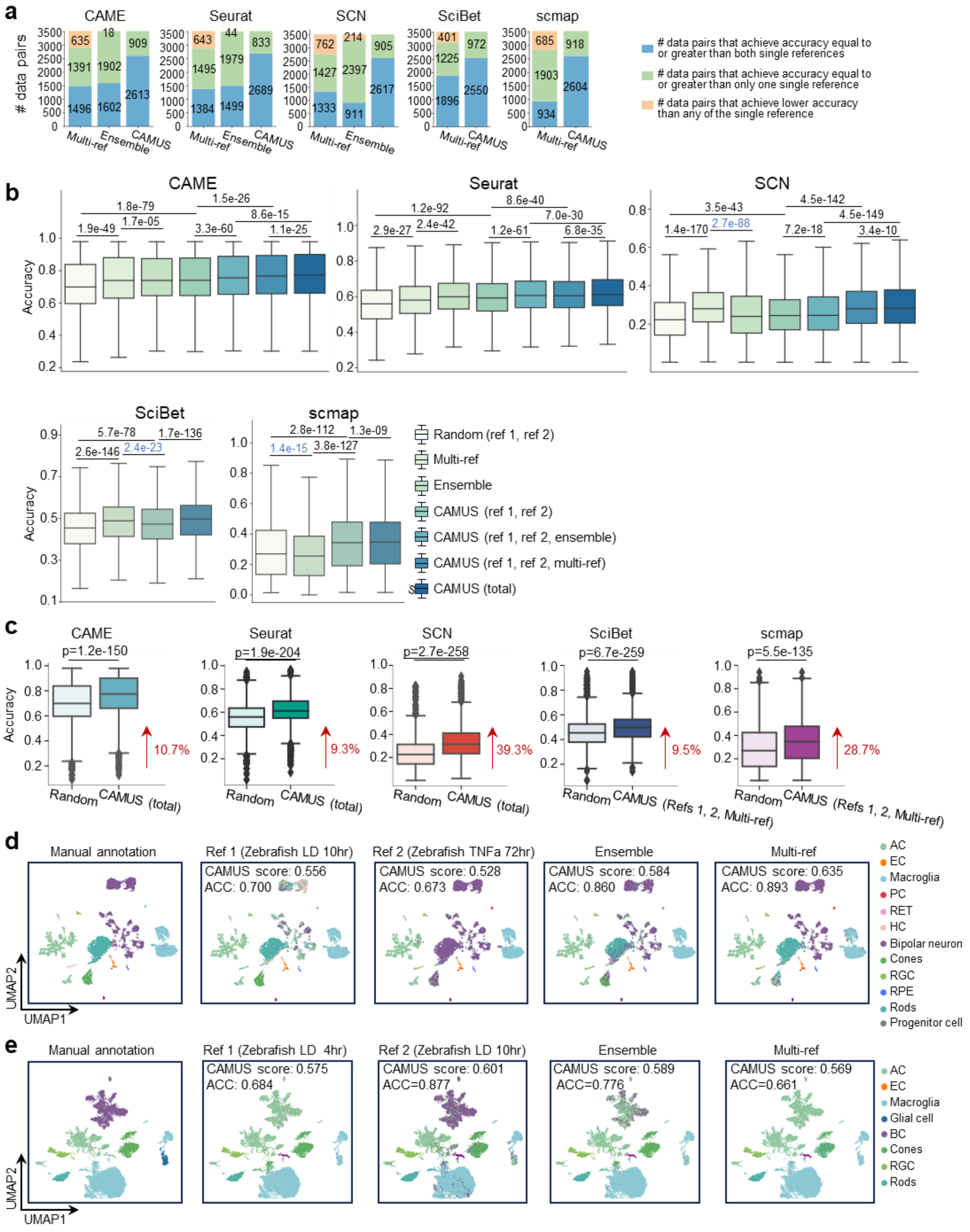


**Supplemental Figure S5. a**, Stacked bar plot showing the improvement of various reference integration/selection strategies over a single reference. **b**, Boxplots showing the annotation accuracy for: random selection between two references (ref1, ref2); multi-reference; ensemble; and CAMUS-based selection among (i) ref1, ref2, (ii) ref1, ref2, ensemble, (iii) ref1, ref2, multi-reference}, and (iv) ref1, ref2, ensemble, multi-reference (“total”). P-values are from paired t-tests; blue indicates the right-hand group has a lower mean than the left, black indicates the opposite. **c**, Boxplot indicating the accuracy of cell type annotations. The references are selected randomly or those selected through CAMUS after applying various reference integration strategies. The p-value was calculated using the paired t-test. The data following the red arrow represents the percentage improvement for the median value relative to random selection. For the random selection, we performed five independent replicates for each setting and report the average values. **d**, UMAP plot of the mouse NMDA P60 dataset from the retina. Cells are colored by annotation of CAME, where using single reference zebrafish NMDA 20hr (the first panel), zebrafish TNFa 36hr (the second panel), the ensemble annotation results (the third panel), and the multi-ref annotation results (the fourth panel). **e**, UMAP plot of the chick p10 dataset from the retina, cells are colored by annotation of CAME, where using single reference zebrafish LD 4hr (the first panel), zebrafish LD 10hr (the second panel), the ensemble annotation results, the multi-ref annotation results (the fourth panel).

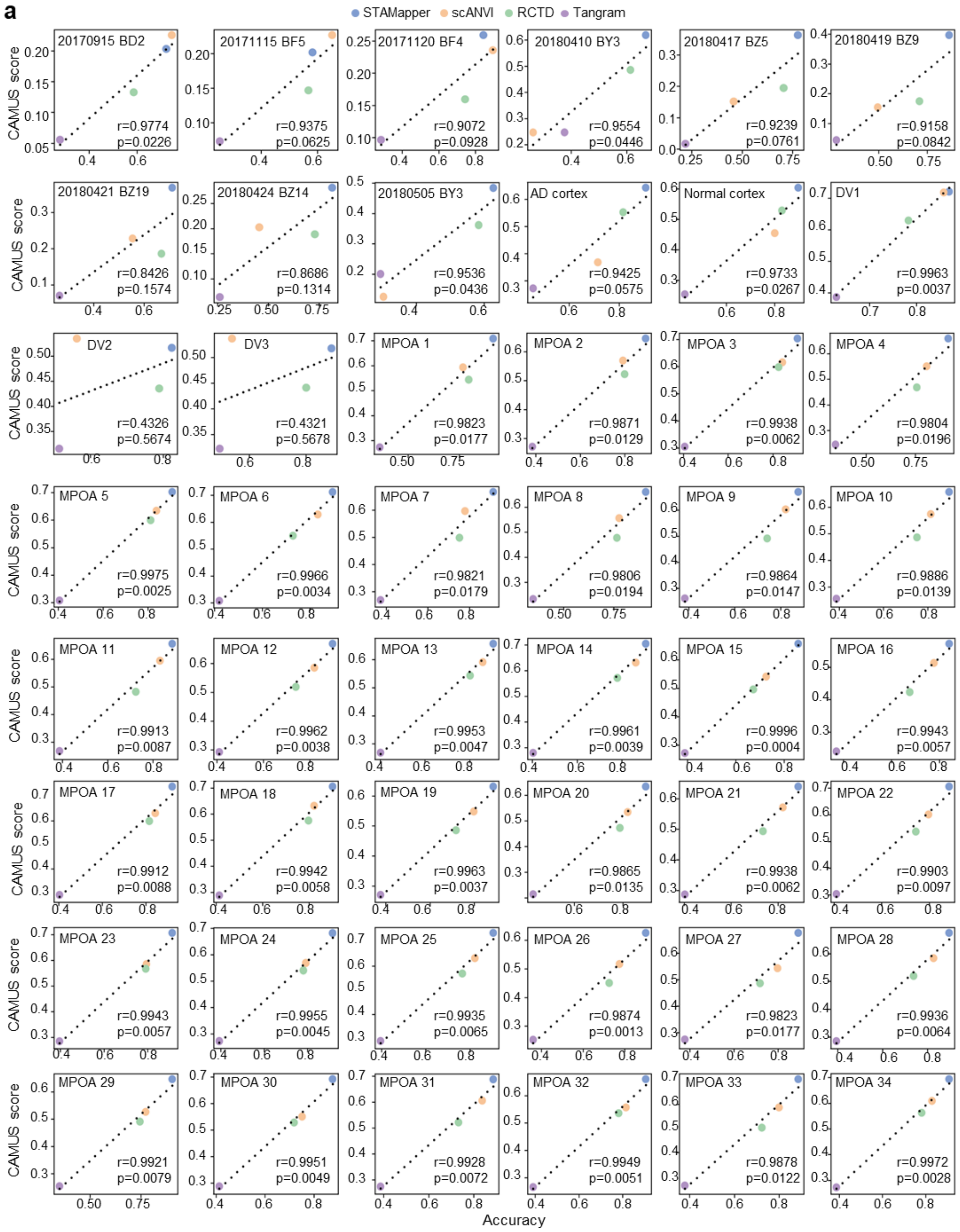


**Supplemental Figure S6 (continue)**

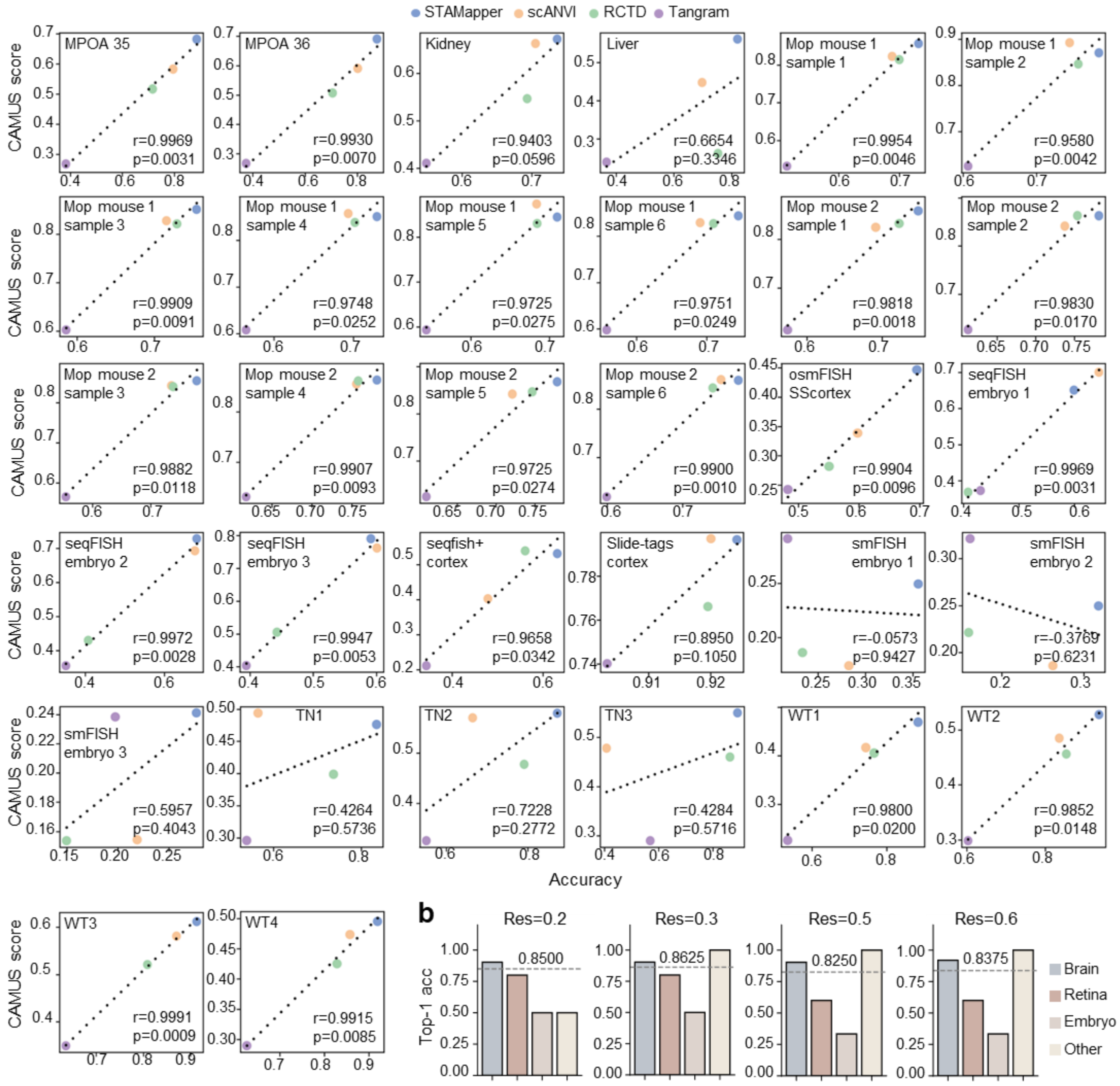

**Supplemental Figure S6. a**, Scatter plot showing the correlation between the CAMUS score and the annotation accuracy across different methods, in 80 scST data pairs. Medial preoptic area (MPOA). **b**, Bar plot showing the top-1, top-2, and top-3 accuracy of selecting the best methods by CAMUS under different resolutions for annotating scST datasets.

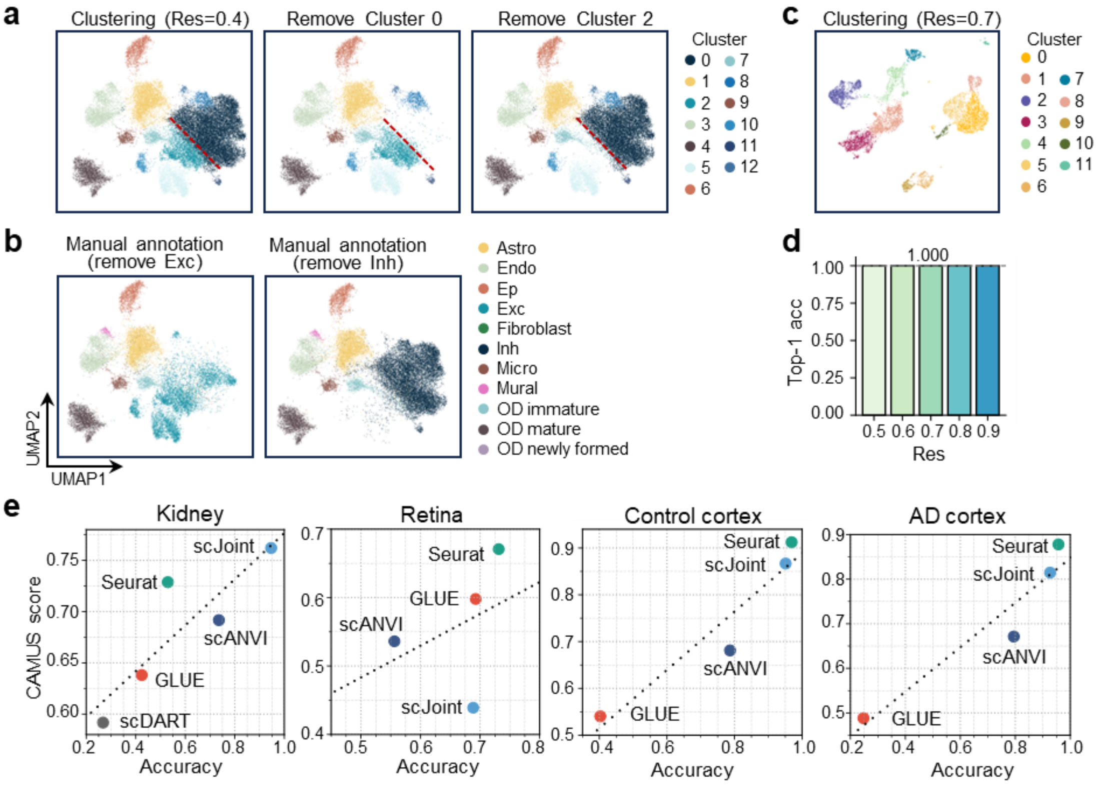


**Supplemental Figure S7. a**, UMAP plots of the mouse hypothalamic region scST dataset from the retina, cells are colored by clustering results. We removed cells from cluster 0 for the middle panel and cluster 2 for the right panel. **b**, UMAP plots of the mouse hypothalamic region scST dataset from the retina. Cells are colored by manual annotation. We removed the excitatory neurons for the left panel and the inhibitory neurons for the right panel. **c**, UMAP plots of the PBMC scATAC-seq dataset. Cells are colored by clustering results. **d**, Bar plot showing the top-1 accuracy for selecting the best methods by CAMUS under different resolutions for annotating scATAC-seq datasets. **e**, Dot plot showing the positive relationship between CAMUS score and the annotation accuracy. AD, Alzheimer's Disease. We showed the results of scDART for the Retina, Control cortex, and AD cortex data due to its runtime exceeding a week.

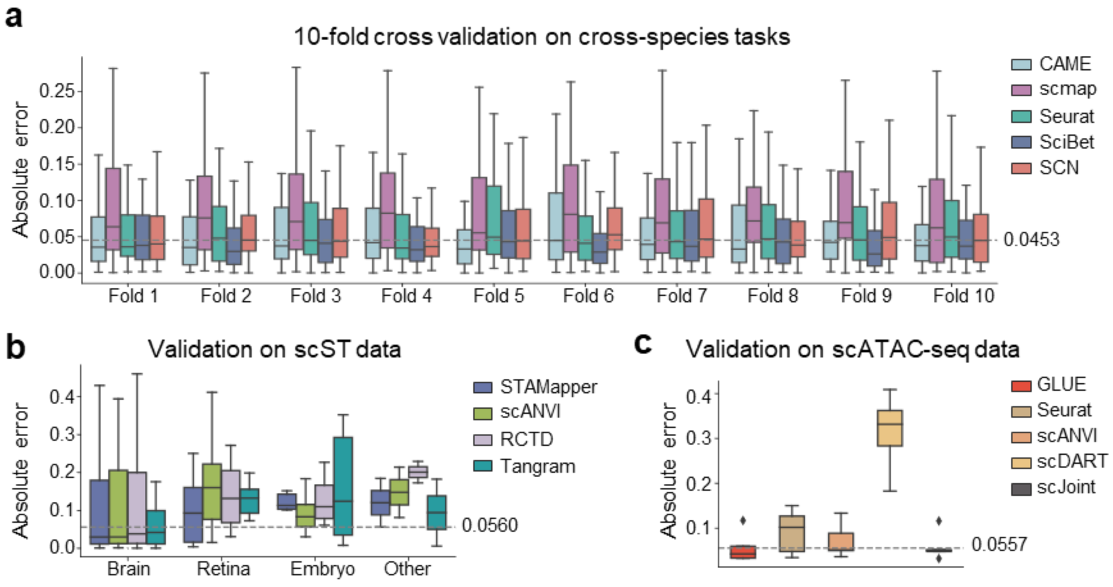


**Supplemental Figure S8. a**, 10-fold cross-validation on 672 pairs of cross-species datasets using CAMUS and other features to estimate the annotation accuracy of five different methods. The dashed line shows the median value of absolute error. **b**, Validation on 80 scST datasets, using the model trained on 672 pairs of cross-species datasets annotated by CAME, Seurat, and SciBet, respectively. **c**, Validation on five scATAC-seq datasets presented in Figure 4C using the model trained on 672 pairs of cross-species datasets annotated by CAME, Seurat, and SciBet, respectively.
